# Complex regulatory interactions at *GDF5* shape joint morphology and osteoarthritis disease risk

**DOI:** 10.1101/2024.11.01.621374

**Authors:** Clarissa R. Coveney, David Maridas, Hao Chen, Pushpanathan Muthuirulan, Zun Liu, Evelyn Jagoda, Siddharth Yarlagadda, Mohammadreza Movahhedi, Benedikt Proffen, Vicki Rosen, Ata M. Kiapour, Terence D. Capellini

## Abstract

Our ability to pinpoint causal variants using GWAS is dependent on understanding the dynamic epigenomic and epistatic context of each associated locus. Being the best studied skeletal locus, *GDF5* associates with many diseases and has a complex cis-regulatory architecture. We interrogate *GDF5* regulatory interactions and model disease variants *in vitro* and *in vivo*. For all regulatory regions we see that local epigenetic activation/repression impacts patterns of joint-specific expression and disease risk. By modeling the most cited risk variant in mice we found that it had no impact on expression, joint morphology, or disease. Yet, we identified significant epistatic expression interactions between this risk variant and others lying within regulatory regions subject to repression or activation. These findings are important lessons on how regulatory interactions and local epistasis work in the etiology of disease risk, and that assessment of individual variants of high GWAS significance need not alone be considered causal.

**Teaser:** Genetic interactions at the most studied skeletal disease locus reveal hidden complexities in pinpointing causal mutations.

## Introduction

Osteoarthritis (OA) is a serious aging disease and the leading cause of disability worldwide, affecting ∼7.6% of the global population (1). Knee and hip OA are highly heritable (40-60%) and for both joints, morphology and mechanical injury are additional risk factors (2, 3, 4, 5, 6). Indeed, ∼50% of hip OA cases result from undiagnosed developmental dysplasia of the hip (DDH) (7, 8) while recently, knee shape has been linked to osteoarthritic disease (5, 9). For both joints, developmental genetic programs underlying cartilage formation prior to and during endochondral ossification aid in determining joint shape (10). The *Growth Differentiation Factor Five* (*GDF5*) gene, expressed during embryonic joint formation (11, 12, 13), is critical in this regard; its loss in the *brachypodism* mouse model (*bp/bp*) results in serious knee and hip malformations (14). When challenged with collagenase, *bp/bp* mice go on to develop OA (15).

Of the numerous OA Genome-wide Association Study (GWAS) loci, *GDF5* is also the most replicable. Hip/knee OA and DDH GWAS consistently reveal ∼100 variants on a common 130 kb risk haplotype spanning the *GDF5* regulatory locus. We previously whittled down this haplotype to two causal variants for knee OA and DHH, revealing their unique impacts on joint shape and disease. We first created a mouse model harboring a knee OA risk variant, rs6060369, in the downstream *GDF5* regulatory region (enhancer/repressor), *R4*, and saw it caused statistically significant morphological changes to femoral condyles and tibial plateaus, and a 30% increase in knee OA risk(9). This finding mirrored the morphological changes observed in OA patients harboring the risk allele. We then generated a mouse model, harboring a different risk variant, rs4911178, present in the *GDF5* growth-plate enhancer, *GROW1*. This variant resulted in alterations to acetabular and femoral neck shape but not the knee proper; with DDH patients stratified for this risk allele showing the same directions of effect (9). Despite this work, on this haplotype there still exist the most associated risk variants, rs143383 and rs143384 (16, 17, 18, 19, 20, 21), located in *5’UTR* of *GDF5*. While reporter gene studies (22) and allele-specific expression (ASE) studies on patient knee tissue (23) have functionally tested these variants, their functional interrogation *in vivo* is lacking, and thus it is unclear if they are relevant to joint disease.

Importantly, ours and others research (14, 18, 19, 21, 24, 25, 26) have led to the *GDF5* regulatory locus as one of the most well-annotated loci in the genome. Besides *R4* and *GROW1*, four other identified regulatory regions (enhancers/repressors) (Figure 1) reside upstream or downstream of *GDF5*. All six enhancers drive different *GDF5* expression patterns *in vivo* as assessed via *lacZ* mouse transgenesis (at embryonic day (E)14.5) and epigenomic studies on developing (E50’s-E60’s) human joints; findings which demonstrate that the mouse and human regulatory loci are functionally orthologous. Yet, as extensive as this work is, a major issue is that no regulatory region alone is able to reproduce the entire endogenous expression pattern (of *GDF5*), indicating that regulatory regions must work together to either enhance, repress, or restrict expression in a modular and tissue specific fashion. Elucidating this regulatory activity for *GDF5*, and how locus-specific activation and repression mechanisms operate, should give us insights into how disease risk variants function. Here, we set out to further explicate and characterize the interactions of *GDF5* regulatory regions, interrogate the function of the leading OA GWAS risk variants rs143383/rs143384, and explore potential variant interactions across the *GDF5* locus.

**Figure 1.**
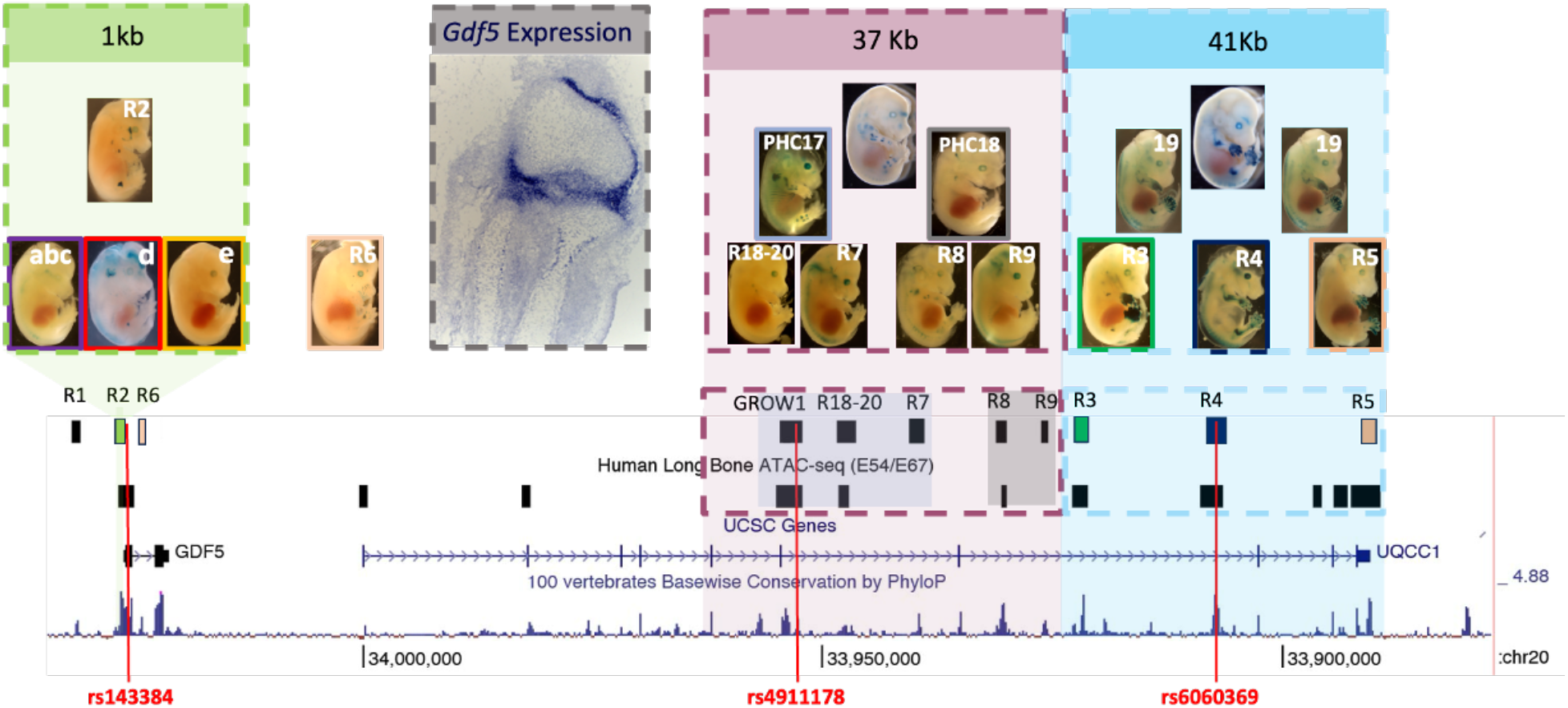
Depiction of the upstream and downstream regulatory regions of *GDF5*. Modified UCSC Genome browser view (hg19) depicting *GDF5* and *UQCC1* genes accompanied by Human Long Bone (proximal tibia and femur, distal femur) ATAC-Seq data (E54/E67), *GDF5* regulatory elements (*R1-R9, R18-20, GROW1*), followed by UCSC Gene locations and peaks of PhyloP100ways conservation. Locations of three variants, rs143384 in the *5’UTR* of *GDF5* rs4911178 in *GROW1*, and rs6060369 in *R4*, overlap with regulatory sequences in embryonic human tissues and mouse are indicated in red. Above, images of transgenic embryos collected at E14.5 depict location of expression for each regulatory element tested including an upstream 1kb region encompassing the *R2* regulatory region (green), a downstream 37kb (purple) region encompassing *GROW1, R18-20, R7, R8* and *R9* and a 41kb (blue) region encompassing *R3, R4* and *R5*. Endogenous *Gdf5* expression in histological tissue of the embryonic knee joint, E14.5 (grey).

## Materials & Methods

### Ethics

All experiments performed on adult or embryonic mice, including euthanasia, have been approved by Stanford University and Harvard University’s respective Institutional Animal Care and Use Committees (IACUC) with protocols (SU, 10665; HU, 13-04-161-2). No human subjects were used. As we are exploring the functional effects of previously published OA GWAS variants, such experiments are not performed in living humans, so no patients were used or involved in this study.

### Animal models

The *5’UTR*^rs143384-T/ rs143384-+^ single allelic replacement mouse line contains a single “T” base-pair replacement of the orthologous human rs143384 variant in the *5’UTR* at mm10 position (chr2:155,945,103-155,945,103). The *R2de* mouse model contains a specific 428bp deletion of the *R2d+e* regulatory region plus adjacent sequence at mm10 position (chr2:155,945,327-155,945,755). Both were CRISPR-Cas9 generated on C57BL/6J *Mus musculus* backgrounds by Applied StemCell as previously described(9).

### Transgenic mice

Transgenic mice were generated as previously described(19). Briefly, transgenic mice were generated by pronuclear injection into FVB or C57BL6/CBA F1 fertilized oocytes (27, 28). Embryos were collected at E14.5 for X-gal staining. For each construct, multiple transgenic embryos derived from independent integration events were analyzed and only consistent patterns are reported (Supplementary Table 1).

### X-gal staining/in situ hybridization

Whole mount β-galactosidase activity was performed as described with minor modifications (28). Embryos were fixed in ice cold 4% PFA in PBS, hemisected then fixed for an additional 15 minutes at 4°C. Following 3 washes in wash buffer, embryos were stained for 16-24 hours in the dark with 1 mg/ml X-gal in staining buffer at room temperature. Embryos were then washed and fixed in 4% PFA for 5 hours. For sectioning, X-gal stained embryos were placed in sucrose solution before being embedded in gelatin, then cryosectioned at 25um. Nuclear Fast Red was used to counterstain sections. *In situ* hybridization: Antisense and sense digoxigenin-labeled probes for *in situ* hybridization were generated for *Gdf5* (29).

### BARX binding site analysis

To find upstream transcription factors predicted to bind to *R4* regulatory sequences, we used UNIPROBE (30, 31). At the recommended enrichment threshold (0.4), UNIPROBE identified over 1,000 sites (i.e., specific 8-mer sequences bound by a transcription factor) in mouse/human *R4* sequences. Predicted factors were intersected with expression/phenotypic data (Eurexpress, http://www.eurexpress.org/ee/; Genepaint, http://www.genepaint.org/Frameset.html; Mouse Genome Informatics, http://www.informatics.jax.org) to narrow down to factors expressed or required in limbs and joints; *Barx1-2* displayed overlap with known *Gdf5* expression patterns, specifically at gestational days when *R4* enhancer was active (32, 33, 34). We next engineered *R4* sequences carrying site-specific BARX mutations and these were synthesized by GenScript. Supplementary document shows the sequence changes per BARX site as calculated using UNIPROBE enrichment score analysis. This technique allowed us to identify those sequences where BARX could bind strongly (i.e. wild type, as above) versus those where BARX could no long bind (mutated sites). Each wild-type (WT) or mutated *R4* element was cloned between *R3* and *R5* within the Hsp68 *lacZ* reporter (Fig. 3b). Each resulting concatenated construct was then used to generate multiple independent E14.5 transgenic mouse embryos for *lacZ* expression analysis (Supplementary Table 1).

### Allele-specific expression

Pyrosequencing was performed to calculate the allelic ratio of C57BL/6J *R2de (*or *5’UTR* rs143384 mouse ‘T’ allele) in the heterozygous state to 129×1/SVJ (wild-type). Heterozygous ratios determined from cDNA products were then normalized by the ratio of wild-type C57BL/6J to 129×1/SVJ genomic products, amplified from known 1:1 mixtures of each sequence. A non-parametric permutation test was used to assess the significance of affect between the wild-type and heterozygous allelic expression allele in R.

### Histomorphology

The left hind limbs of a minimum of 4 animals/genotype/sex/time point were obtained for histological analysis to assess cartilage integrity. Post sacrifice, skin and excess muscle was removed, and each limb was fixed in 10% neutral buffered formalin for 24 hours at room temperature before being decalcified in 14% Ethylene-diaminetetraacetic acid (EDTA) at pH 7.5 for 8 days. Formalin-fixed tissues were sent to the Massachusetts General Hospital Center for Skeletal Research (CSR) Histology & Histomorphometry Services core facility and processed in one batch for proper sectioning and histological staining using Fast Green and Safranin O (Saf O) staining. Embedding, sectioning, and staining were performed without knowledge of genotype. A minimum of 10 coronal sections taken throughout the joint (60-80um levels) were generated per knee joint. Sections were blind scored by two readers (CC and AK/BP) using the summed Osteoarthritis Research Society International (OARSI) scoring method of OA in the mouse. Briefly, per slide, each quadrant (medial and lateral tibial plateaus, medial and lateral femoral condyles), was assigned a score and summed, with 0 representing intact cartilage, and 6 representing cartilage erosion down to the bone extending >75% of the articular surface. The top 3 summed slides per joint were added to generate one final score per joint. One way ANOVA (*R2de*) was used to compare OARSI scores between genotypes whereas, Two-way ANOVA (*5’UTR*^rs143384/rs143384^) was used to compare OARSI scores between the sex and genotype.

### Functional Analysis of *GROW1*/*R4* + *5’ UTR* risk and non-risk variants using luciferase reporter assays

CHON-002 cells, derived from human fetal femoral growth plate chondrocytes at 18 weeks of gestation (female), were sourced from ATCC (CRL-2847), NIH/3T3 (Hoekstra Lab, Harvard University) cells and T/C28a2 cells were cultured at 37°C in a 5% CO_2_ environment using ATCC’s complete growth medium, which consists of Dulbecco’s Modified Eagle’s Medium (DMEM) supplemented with 10% fetal bovine serum (FBS), 50 μg/mL penicillin-streptomycin, and 0.1 mg/mL G-418.

For transfection experiments, CHON-002 (*GROW1* constructs) cells, T/C-28a2 (*R4* constructs and *R4* single variant constructs previously described(26)) or NIH/3T3 (*R4* single variant constructs) cells were seeded in 96-well plates at a density of 1–4 × 10^4^ cells per well and cultured in DMEM supplemented with 10% FBS for 24 hours. Transient transfections were performed using Lipofectamine™ 3000 (Invitrogen L3000015) in serum-free DMEM, according to the manufacturer’s protocol. The transfection complex was prepared by diluting Lipofectamine 3000 in Opti-MEM Medium (5 μL Opti-MEM + 0.3 μL Lipofectamine 3000 per well of a 96-well plate). Separately, a DNA-P3000 mix was prepared (5 μL Opti-MEM + 100 ng (or 200ng for NIH/3T3) DNA + 0.2 μL P3000 Reagent per well). The diluted DNA solution was then added to the diluted Lipofectamine 3000 in a 1:1 ratio and incubated at room temperature for 10-15 minutes. Cells were transfected with a firefly luciferase reporter vector containing a *GROW1* and *5’UTR* or a *R4* and *5’UTR* and fusion sequence with various combinations of *GROW1* (rs4911178) and *5’UTR* (rs143383 and rs143384) variants and *R4* (rs6060369) and *5’UTR* (rs143383 and rs143384) (100 ng total). An empty pGL4.23 luciferase vector (100 ng) was used as a control. To normalize transfection efficiency, the pRL-CMV Renilla luciferase vector (Promega; E226A) was co-transfected. The epistatic activity of the fusion variant combinations was measured 24 hours post-transfection using the Dual-Luciferase Reporter Assay System (Promega, TM040) on an Agilent BioTek Synergy Neo2 Hybrid Multimode Reader (Thermo Fisher Scientific, USA), following the manufacturer’s instructions. Standard and variant synthesis of either *R4* or *GROW1* sequences concatenated to the *5’UTR* including added 5’ KpnI and 3’ HindIII sequences with sequence verification and custom cloning into pGL4.23 custom (Ampicillin) via 5’ KpnI and 3’ HindIII (by recombination) were generated by Genewiz and delivered as a mini-scale DNA sample (see Supplementary Materials for sequences of each construct). The constructs used for the functional analysis of *GROW1* and *5’UTR* variants in the Dual-Luciferase Reporter Assay are shown in Supplementary Materials.

### CRISPR methods

CRISPR targeting of regulatory regions *R9*, and *R2de in vitro*. All sgRNAs flanking (1) the *R9* regulatory region (2) and the *R2de* region were designed using MIT CRISPR Tools (http://crispr.mit.edu) and synthesized by Integrated DNA Technologies, Inc (Coralville, Iowa), and cloned into a PX458 vector as previously described in published protocols(35). See supplementary materials for sequences and chromosomal locations of sgRNAs. All guide RNAs, were tested for deletion efficiency of respective human elements in cultured T/C-28a2 cells (n=3 biological replicates per assay). T/C-28a2 cells were maintained as described above and seeded in a 6 well plate 24 hours before transfection. Transfection efficiency was measured using a fluorescence microscope (>70% of cells were GFP positive). Extraction of DNA was performed using the E.Z.N.A Tissue DNA Kit (Omega Bio-Tek, Norcross, GA), and respective regulatory elements were amplified using PCR with primers flanking each sgRNA location (Supplementary Table 2), followed by purification from 1% agarose gel (E.Z.N.A Gel Extraction Kit). Sanger sequencing was used to verify each successful targeting event

Impacts of each modification on *GDF5, UQCC*, and *CEP250* expression were assessed by extracting RNA from control and CRISPR-Cas9 targeted T/C-28a2 or NIH/3T3 cells (n = 3 biological replicates, with three technical replicates per experiment per condition) using Trizol Reagent (Thermo Fisher Scientific, Springfield Township, New Jersey) and Direct-zol™ RNA Miniprep kit (ZYMO). SuperScript III First-Strand Synthesis System (Thermo Fisher Scientific) was used to prepare cDNA. qRT-PCR analysis was then performed with gene specific primers (9, 26) and Applied Biosystems Power SYBR master mix (Thermo Fisher Scientific) with GAPDH house-keeping gene as an internal control.

### Micro-CT and anatomical measurements

Acetabular joint, femur, and tibia from right hind limbs were collected using high-resolution Micro-Computed Tomography (μCT40, SCANCO Medical AG, Brüttisellen, Switzerland). Scan parameters were 12 μm3 isotropic voxel size, 70 kVp peak X-ray tube intensity, 114 mA X-ray tube current, and 200 ms integration time. Resultant DICOM images were exported for measurements following anatomical features in Osirix MD v7.5 (Pixemo SARL, Bernex, Switzerland). First, using previously described methods (9), measurements were taken of the acetabulum (depth, diameter, inclination), proximal femur (valgus cut angle, neck-shaft angle, neck length, neck diameter, head offset, head diameter), distal femur (bicondylar width, notch width, condylar width (medial and lateral), condylar curvature (medial and lateral), trochlear width (medial, central, and lateral), trochlear groove depth, trochlear angle), and proximal tibia (plateau width, posterior tibial slope (medial and lateral), tibial spine height (medial and lateral)). All measurements displayed strong inter- and intra-examiner reliability (ICC > 0.78). Second, MicroCT-DICOM images were segmented to generate 3D models of each bone using image processing software (Mimics v17.0, Materialise). 3D models were imported to 3-matic software package (v9.0, Materialise) and co-registered together using a global n=point registration method. 3D models of wild type and homozygous mice were used to generate 3D heatmaps indicating the geometrical differences between genotypes for each mouse line. The heatmaps were generated by calculating the distance between the corresponding points in co-registered models, where dark blue indicates the maximum deviation in the negative direction and red indicates the maximum deviation in the positive direction.

### Statistical analysis

Expression data for *GDF5, CEP250*, and *UQCC1* were normalized relative to *GAPDH* house-keeping gene expression and compared between control and *R2de* enhancer deletion. All data are presented as the mean ± SEM unless otherwise stated. Individual pairwise comparisons between control and experimental condition were analyzed by two-sample, two-tailed Student’s t-test, with p < 0.05 regarded as significant. N = 4 technical replicates per biological replicate (3 biological replicates).

## Results

### Activation/repression domains within a 37 kb sequence containing *GROW1*

We previously reported on a broad 37kb human sequence downstream of *GDF5* (purple region, Fig. 1, Fig. 2a) that drives *GDF5* expression only in the sub-perichondral region of long-bone growth plates, identical to that of *GROW1, a* 2.54kb enhancer containing a key DDH risk variant (rs4911178). Interestingly, this 37kb sequence consists of a ∼12.17kb subregion *PHC17* (Fig. 1, light grey) that drives expression within the entire growth plate chondrocytes (i.e., not only in the sub-perichondral region) and a ∼10kb subregion *PHC18* (Fig. 1, dark grey) that drives expression in the shoulder joint proper (Fig.2b-c). Since *Gdf5* is not endogenously expressed throughout the growth plate (Fig. 1), this means that built-in or adjacent to *PHC17* there are repressors that restrict expression to the sub-perichondral space. We therefore first tested two conserved *PHC17* sub-regions, one called *R18-20* (∼2.7kb from *GROW1)*, and another called *R7* (∼6.9kb from *GROW1*). We found that neither drive growth plate expression (Fig. 2b-c), but rather at other joint and digit sites. As noted, the adjacent *PHC18* (Fig. 2c) drives expression in the shoulder, which is not a pattern controlled by the larger 37kb sequence. We next tested two conserved regions within *PHC18* termed *R8* and *R9*. Surprisingly, *R8* alone drives expression within many more hind limb and forelimb joints, indicating that it is being repressed in *PHC18* and the 37kb constructs. Upon testing *R9* we found no expression in limbs indicating it might act as a repressor, especially when considered in *PHC18* and 37kb parental constructs. To test this, we deleted *R9* in human T/C-28a2 chondrocytes and observed a significant increase in *GDF5* expression, thus revealing a normal repressor role (Fig. 2d). We note that in this *R9* repressor there exists two OA GWAS risk variants (rs2378349 and rs2248393) that may modulate repressor and thus *GDF5* expression levels. We performed a luciferase reporter test of these two variants in T/C-28a2 chondrocytes and observe some impacts of the risk allele on *GDF5* expression (see Supplementary Figure 1c). Thus, in considering both *PHC17* and *PHC18* within the 37kb region, we note that the endogenous growth plate sub-perichondral expression, recapitulated by both 37kb and *GROW1* regions (Fig. 2a) results from two potential repression systems: (1) a localized system within *PHC17* via *R18-20* and/or *R7* and (2) a long-range system within *PHC18* via the actions of the *R9* repressor.

**Figure. 2.**
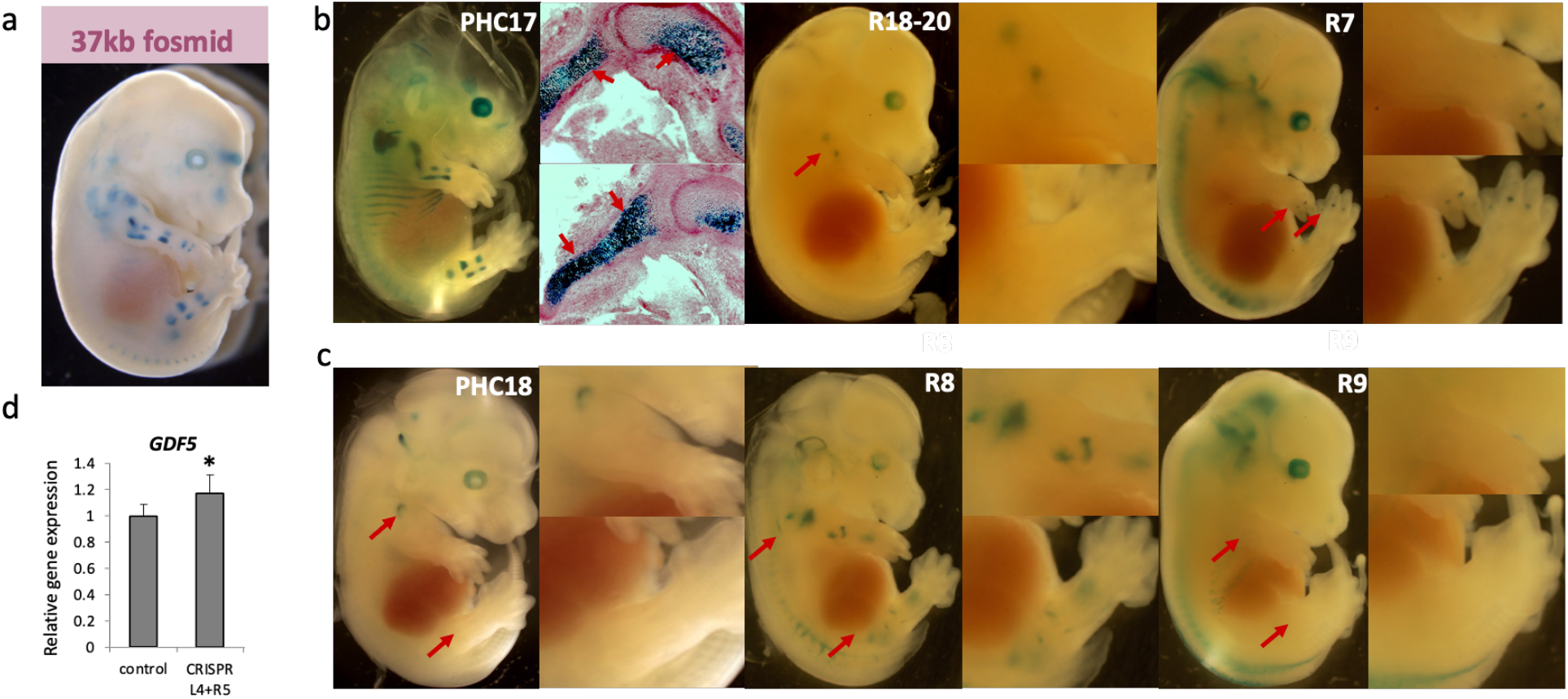
Restriction of *GROW1* to the perichondrium exerted by downstream regulatory regions. Transgenic embryos driving *lacZ* were collected at E14.5 of (a) 37kb fosmid driving growth plate expression and (b) regulatory region *PHC17* (*GROW1, R18-20*, and *R7*) with two histological panels of the hind limb showing *PHC17*-driven LacZ expression throughout the entire growth plate. (c) Transgenic embryos collected at E14.5 of region *PHC18* (which includes *R8* and *R9*). Red arrows indicate locations of differential expression patterns between regulatory elements. (d) Relative *Gdf5* expression following CRISPR knockout of *R9* in T/C-28a2 cells.

**Figure 3.**
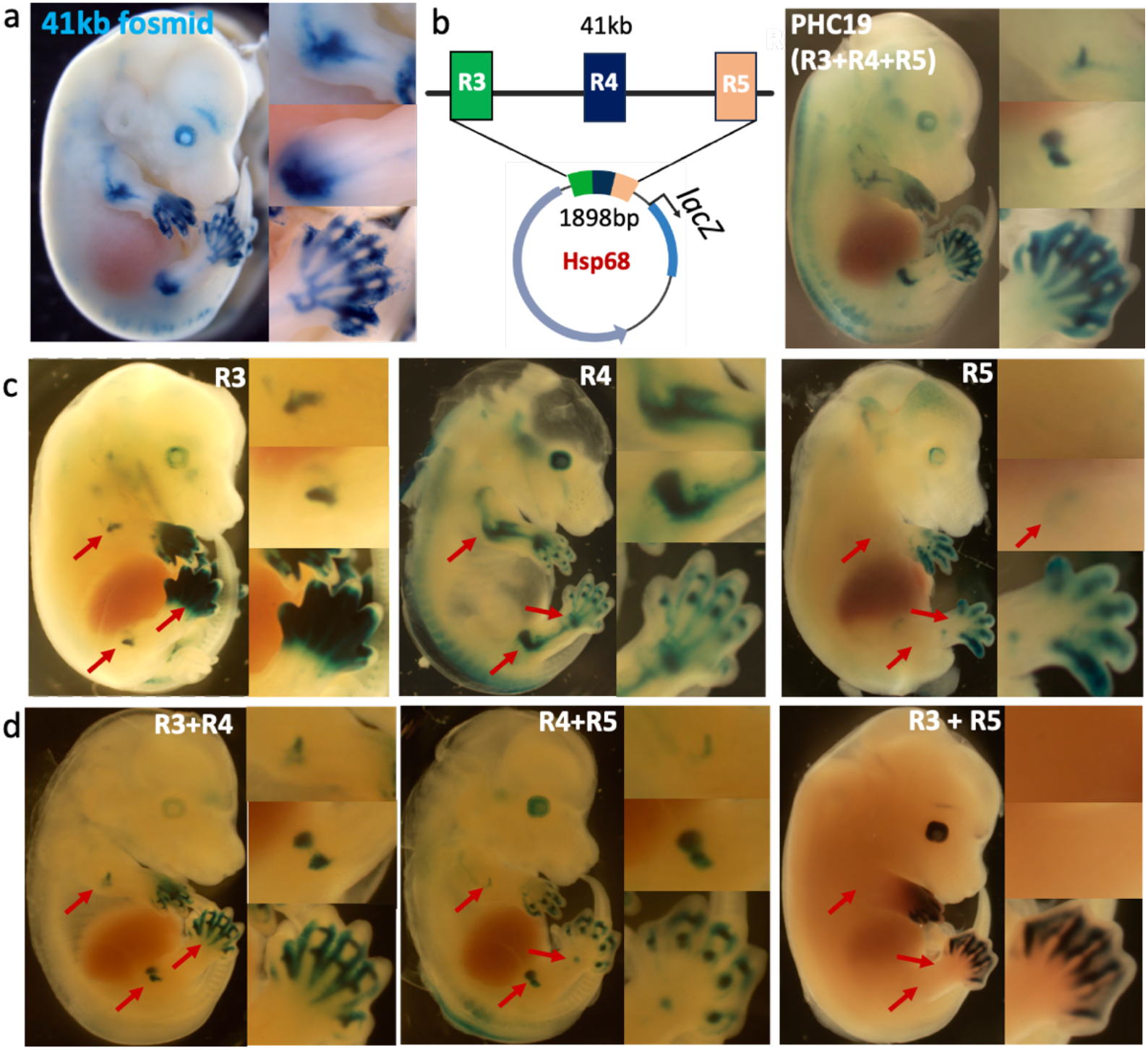
Repressive interactions between nearby regulatory elements affects expression patterns. (a-b) Transgenic embryos collected at E14.5 of (a) the 41kb construct encompassing *R3, R4* and *R5*, and (b) the *PHC19* construct concatenating regulatory elements *R3, R4*, and *R5* (1898bp), both of which drive expression in the same limb domains. Cartoon depiction of construct containing concatenated regions. (c) Transgenic embryos collected at E14.5 of the *R3* region, driving *lacZ* expression in the autopod, and forelimb and hind limb joints, the *R4* region, driving expression in the forelimb, hind limb, and digit synovial joints, and the *R5* region, driving expression predominantly in the digits. (d) Transgenic embryos collected at E14.5 showing how *R3*+*R4* drive expression in the forelimb, hind limb and digit joints with some expression in the digit periphery; *R4*+*R5* drive expression in the forelimb, hind limb and digit joints; and *R3*+*R5* drive expression only in the webbing of the digits. Red arrows indicate areas of expression patterns that change between constructs. See text for details.

### Activation/repression domains within a 41 kb sequence containing *R4*

We had characterized a human 41 kb sequence (blue region, Fig. 1, Fig. 3a), further downstream of *GDF5*, showing it recapitulated the classic *Gdf5* limb joint-specific expression pattern. This regions three-known enhancers drive unique patterns: *R3* in the interdigital space, digital transverse stripes, and knee/elbow joints (*R3*, Fig. 3c); *R4* in the elbow/knee, and digit joints, with weaker expression in hip/shoulder (R4, Fig. 3c); *R5* in the prechondrogenic phalangeal mesenchyme and weakly in elbow/knee (*R5*, Fig. 3c)(19). Alone, *R3, R4*, or *R5* cannot drive the broader 41 kb pattern. Moreover, *R3* and *R5* patterns are not observed by the broader 41 kb sequence, nor are they known *Gdf5* expression territories. Interestingly, when concatenated (in construct *PHC19*, Fig. 3b) *R3*(586bp) + *R4*(975bp) + *R5*(337bp) (i.e.,1898bp out of the 41kb) strikingly generated the 41kb pattern. To understand this complex interaction further, we first concatenated *R3*+*R4* and found together they drive strong expression in the knee/elbow, and digit joints including transverse stripes and some interdigital webbing (Fig. 3d). Here, *R3* acts to restrict knee, elbow, and shoulder expression of *R4*, whilst *R4* represses *R3* expression in the interdigital webbing. We next tested *R4*+*R5* and found together they drive weak expression in the elbow, but strong expression in the knee, transverse stripes of the digits, and weak expression of the metacarpals (Fig. 3d). Here, *R4* acts to suppress *R5* mesenchyme expression whilst *R5* acts to restrict (and weaken) *R4* expression in large joints. Finally, we tested *R3*+*R5* and found together they cause the strong digital mesenchyme expression of *R5* to be lost, while *R3* expression becomes further restricted in the interdigital webbing (Fig. 3d). These studies reveal that in addition to enhancer activities, each regulatory element can act as a repressor.

As we have shown, the ability of a regulatory region to restrict expression appears to be joint and tissue dependent, emphasizing the importance of location for gene regulatory activity to recapitulate endogenous *Gdf5* expression. To emphasize the importance of this finding to causal disease biology, we tested the previously reported *R4* “T” OA risk allele at rs6060369 in two contexts. First, by testing the risk “T” allele (compared to the non-risk “C” allele) in human T/C-28a2 chondrocytes (a cell type in the developing knee), we find that the *R4* element acts as an enhancer, and the “T” risk allele decreases its activity (Supplementary Fig. 1a). However, when the “T” risk allele is tested in a fibroblast line (NIH/3T3 cells; another cell-type in the developing knee), we find it serves to derepress the activity (and thus increase expression towards baseline) (Supplementary Fig. 1b). These findings reveal that as in our *in vivo* LacZ experiments, *R4* in different human cellular contexts can repress or activate, and importantly that its risk variant effects depend on the epigenomic context.

### Predicted transcription factor binding sites in *R4* are required for expression in the forelimb and hind limb joints

While variant roles in disease biology are dependent on the *cis*-regulatory, epigenomic context for which they are located, the transcription factor (TF) or *trans*-environment is also important to consider. We next deeply interrogated the *R4* regulatory element given its importance to knee OA risk. Using published methods (Methods) we identified BARX, a homeodomain protein with known roles in chondrogenesis, as predicted to strongly bind to five locations across *R4* (Fig. 4a)(32). BARX1 and BARX2 are known to be expressed across developing joints, exhibiting strong overlap with *GDF5* expression (32, 33, 34, 36). We next sought to experimentally test each BARX binding site using *in vivo* lacZ approaches (Methods). We first tested all 5 sites within *R4* by mutating each to destroy (through reshuffling) their predicted binding sequence. When all 5 sites were mutated (MUTB1-MUTB5,) *lacZ* expression was restricted only to the digits, reducing expression in digit webbing and eliminating large limb joint expression (Fig. 4 b and 4c and Supplementary Figure 2a and 2b). We next generated separate lacZ constructs with each TF binding site mutated independently and found no impact on expression patterns as a result of MUTB1, MUTB4 or MUTB5 (Fig. 4d, and Supplementary Figure 2c), but strong repression of knee and shoulder expression as a result of MUTB3 (Fig. 4e). MUTB2 (Supplementary Figure 2c) also reduced expression within the limb joints but not as strongly as MUTB3. Interestingly, the OA GWAS associated *R4* variant (rs6060369)(9) lies between MUTB2 (92bp away) and MUTB3 (66bp away), respectively. Overall, these data reveal the importance of transcription factor functioning on *R4* activity and that individual TF binding sites (and variants nearby) can have sizable impacts of the regulatory activity of important enhancers.

**Figure 4.**
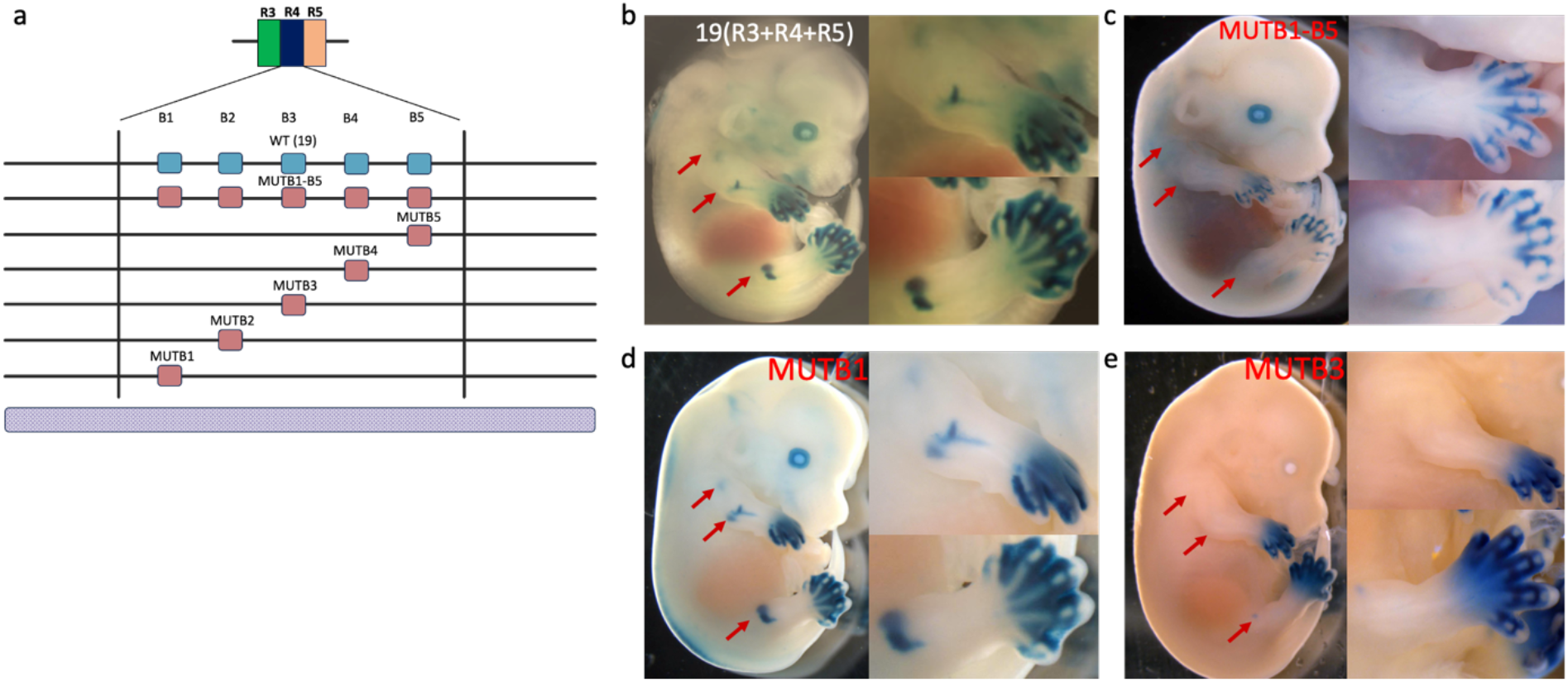
Predicted BARX binding sites are required for *R4* hind limb and forelimb joint expression. (a) Five predicted (BARX) homeodomain binding sites were identified within *R4* and mutated at each individual site, or all were mutated simultaneously. (b) WT transgenic E14.5 embryos of construct *PHC19* (R3+R4+R5) driving expression of *lacZ*. (c) Transgenic E14.5 embryos of construct *PHC19* with all 5 BARX sites mutated, showing loss of expression in the forelimb and hind limb joints. (d) Transgenic E14.5 embryos of construct *PHC19* with only one BARX site mutated (MUTB1) where there is no impact on lacZ expression. (e) Transgenic E14.5 embryos of construct *PHC19* with mutated (MUTB3) where there is a complete loss of forelimb and hind limb joint expression. Red arrows indicate areas of expression patterns that change between constructs.

### Activating/repression domains within the upstream *R2* joint enhancer

We had reported (19) a ∼1 kb enhancer, *R2* (green region, Fig 1.), that drives strong expression within limb joints and recapitulates a subset of the entire endogenous *Gdf5* expression pattern. This element resides near the *GDF5* promoter and *5’UTR*. There are 5 highly conserved regions(19) across the *R2* element denoted *a-e* (Fig. 5a). Interestingly, whilst *a, b*, and *c* do not reproduce expression either independently or together (*a+b+c*) (Fig.5b), sub-elements *d* and *e* can recapitulate expression in the hip and knee or shoulder and elbow, respectively (19). We identified a shorter 112bp sequence within *d* that suppresses elbow and metapodial joint expression (Supplementary Figure 3). Most strikingly, by concatenating *a+b+c* with either *d* or *e* (i.e., *a+b+c+d* or *a+b+c+e*), *abc* suppresses expression in both the hind limb and forelimb. This reveals built-in repression domains within individual enhancers. Strikingly, under any combination of sub-element concatenation, digit expression could not be recapitulated, revealing that digit expression requires the full *R2* regulatory element to be intact for expression.

**Figure 5.**
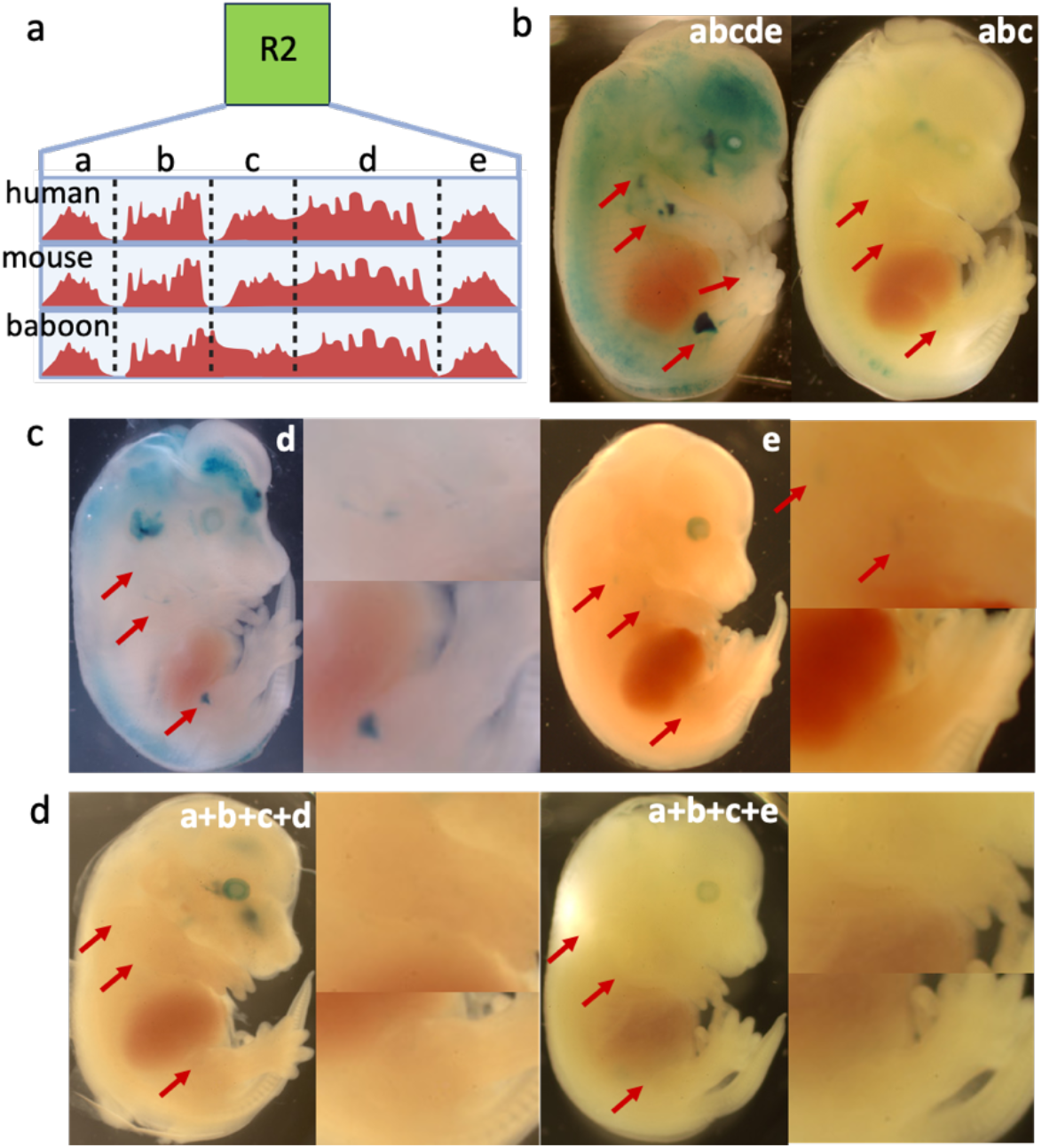
Sub-regions of *R2* repress regulatory region activity. a) Cartoon representation of conservation of 5 sub-regions (a-e) within the *R2* element across different species (19). (b) Transgenic E14.5 embryos of the entire *R2a-e* sequence driving expression in the forelimb, hind limb and digit joints; sub-region *R2abc* is unable to drive expression in any joint, whilst subregions *de* drive expression in hind limb and forelimb joints, but not in digit joints. (c) Transgenic E14.5 embryos show sub-regions *R2d* and *R2e* are able to drive expression in hind limb and forelimb joints respectively. (d) Transgenic E14.5 embryos showing that sub-regions *abc* when placed adjacent to sub-region *d* or *e* represses hind limb or forelimb expression, respectively. Red arrows indicate areas of expression patterns that change between constructs.

### Deletion of *R2de* results in morphological changes not associated with OA development

Enhancer *R2* is adjacent to *GDF5 5’UTR*, in which resides the most associated OA GWAS variant, rs143384. To understand the role of *R2* in large joint (e.g., knee) biology and OA risk, we generated a mouse line with a deletion of the limb-joint-specific *R2de* region. Using allele-specific expression analysis (ASE) on 7 separate E15.5 limb joint sites, we observe a consistent ∼40% decrease in *Gdf5* expression at each site (Supplementary Figure 4a). When deleted in T/C-28a2 human chondrocytes, *GDF5* (but not nearby *UQCC1* and *CEP250*) expression is similarly reduced (Supplementary Figure 4b, 4c and 4d respectively). Given the expression reduction, we performed MicroCT morphological analyses on wild-type (WT), *R2de*^*-/+*^ and *R2de*^*-/-*^ mice at P56 and found statistically significant shape changes across the knee’s femoral and tibial plateau and hip’s acetabulum (Fig. 6a and Supplementary Figure 4f). Strikingly, by 1.5 years, only knee bicondylar width remained statistically impacted (Fig. 6b and Supplementary Figure 4g). By 3D analysis, these changes predominantly locate to the medial femoral condyles at P56 and 1.5years, while at 1.5years hip alterations are ameliorated though some changes remain at the anterior tibial plateau (Fig. 6c). Apart from two WT cases of spontaneous OA at 1.5 years, there are no changes to cartilage integrity measured by OARSI scoring at P56 or 1.5 years of age, and separation of scores by plateau and condyle does not reveal regionally-specific effects (Fig. 6d, 6e and Supplementary Figure. 4e). Likewise, no statistically significant changes are observed in tibial or femoral articular cartilage thickness at either time point (Fig. 6e and Supplementary Figure. 4e). Thus, the loss of *R2de*, which has a marked effect on *Gdf5* gene expression, has no long-term observable impacts on joint morphology and no increased risk of spontaneous OA.

**Figure 6.**
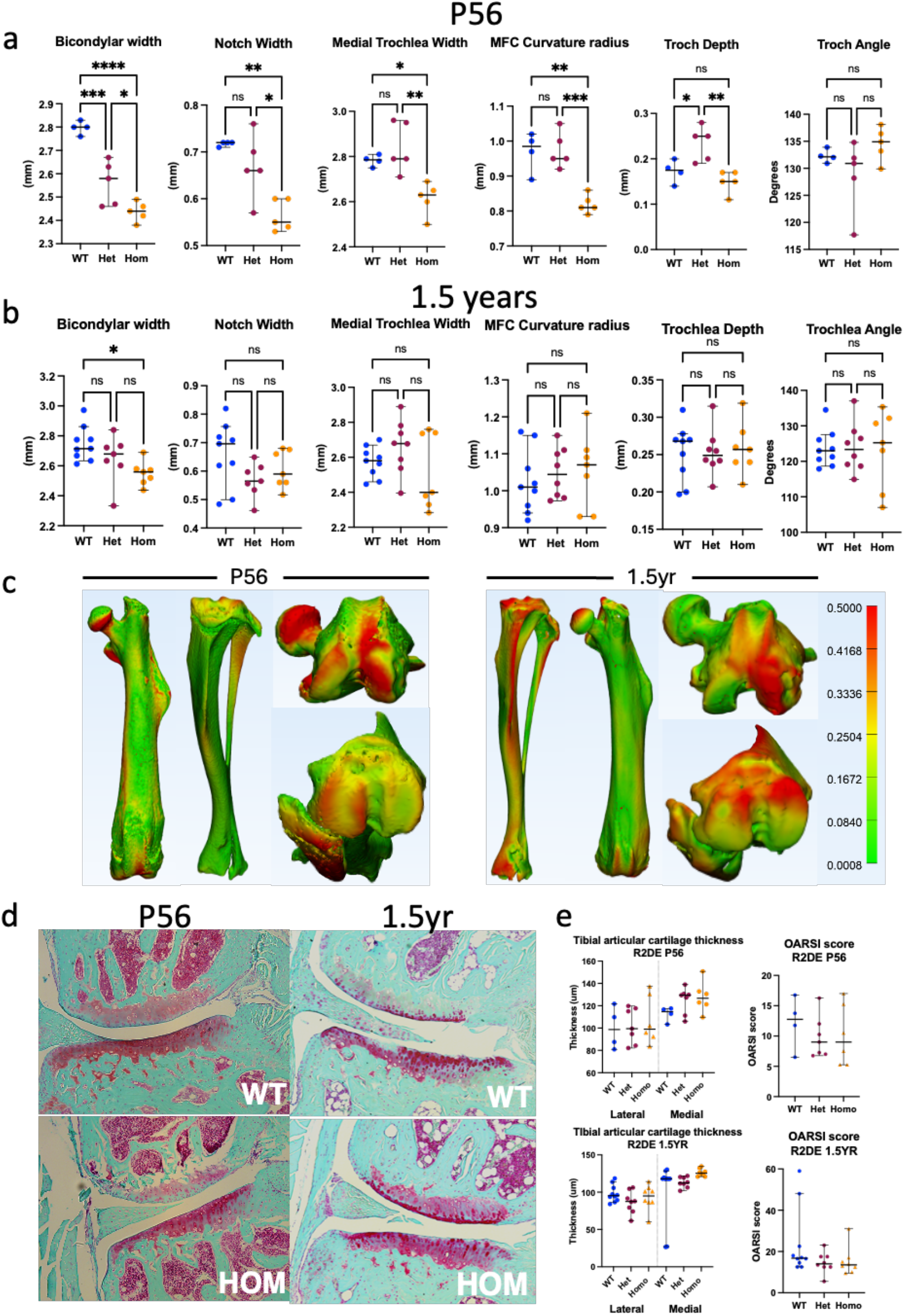
Morphological characterization of the *Gdf5 R2de* enhancer mouse model. (a) MicroCT measurements of significantly different anatomical features in *R2de* deletion mice at P56 (WT n = 4, Het n = 5, Hom n = 6) and 1.5 years (WT n= 8, Het n= 7, Hom n = 7). One-way ANOVA with Tukey-kramer test was used for comparisons between all groups (*p<0.05, **p<0.01, ***p<0.001, ****p<0.0001, bars indicate medians and 95% Confidence Intervals) at both P56 and (b) 1.5 years of age. (c) 3D morphological comparative analysis indicating locations of largest anatomical differences between wild type (WT) and homozygous *R2de* null hind limbs at P56 and 1.5 years. (d) Coronal histological sections of the medial compartment stained with Saf O for representative OA score per timepoint. (e) Tibial articular cartilage thickness measurements of the lateral and medial plateaus at P56 and 1.5 years alongside respective OARSI scores for each time point (P56: WT n = 4, Het = 5 HOMO n = 6; 1.5 years: WT n= 8, Het n= 7, Hom n = 7).

### rs143384 risk allelic mice lack joint alterations and OA disease

Within the *GDF5 5’UTR* is the OA GWAS risk variant, rs143384, a ‘G’ to ‘A’ risk mutation. While others have shown that the variant impacts reporter gene expression *in vitro* or *GDF5* expression in patients (23), no one has tested its role in joint development and homeostasis *in vivo*. We generated a humanized *5’UTR*^*rs143384-A/rs143384-G*^ allelic replacement line. Using ASE at E15.5 on 7 joint sites, we observe no impact of the risk ‘A’ allele on *Gdf5* expression (Supplementary Figure 5a). Using MicroCT imaging on P56 wild-type, *5’UTR*^*rs143384-G/rs143384-A*^, and *5’UTR*^*rs143384-A/rs143384-A*^ mice, we find the “A” allele causes no significant changes to the width and curvature of the knee’s femoral condyles or to the size and slopes of the tibial plateau at P56 (Fig. 7a, 7b, and Supplementary Fig 5c). We observe only a modest impact on notch height in female mice, albeit *GDF5* does not exhibit sex effects on OA risk in any GWAS to date. Histologically, we observe no increase in disease risk between each genotype, or sex by OARSI scoring at P56 (Fig. 7c and Supplementary Figure 5b). We conclude that modeling the variant alone reveals little-to-no impact on knee OA biology and disease risk.

**Figure 7.**
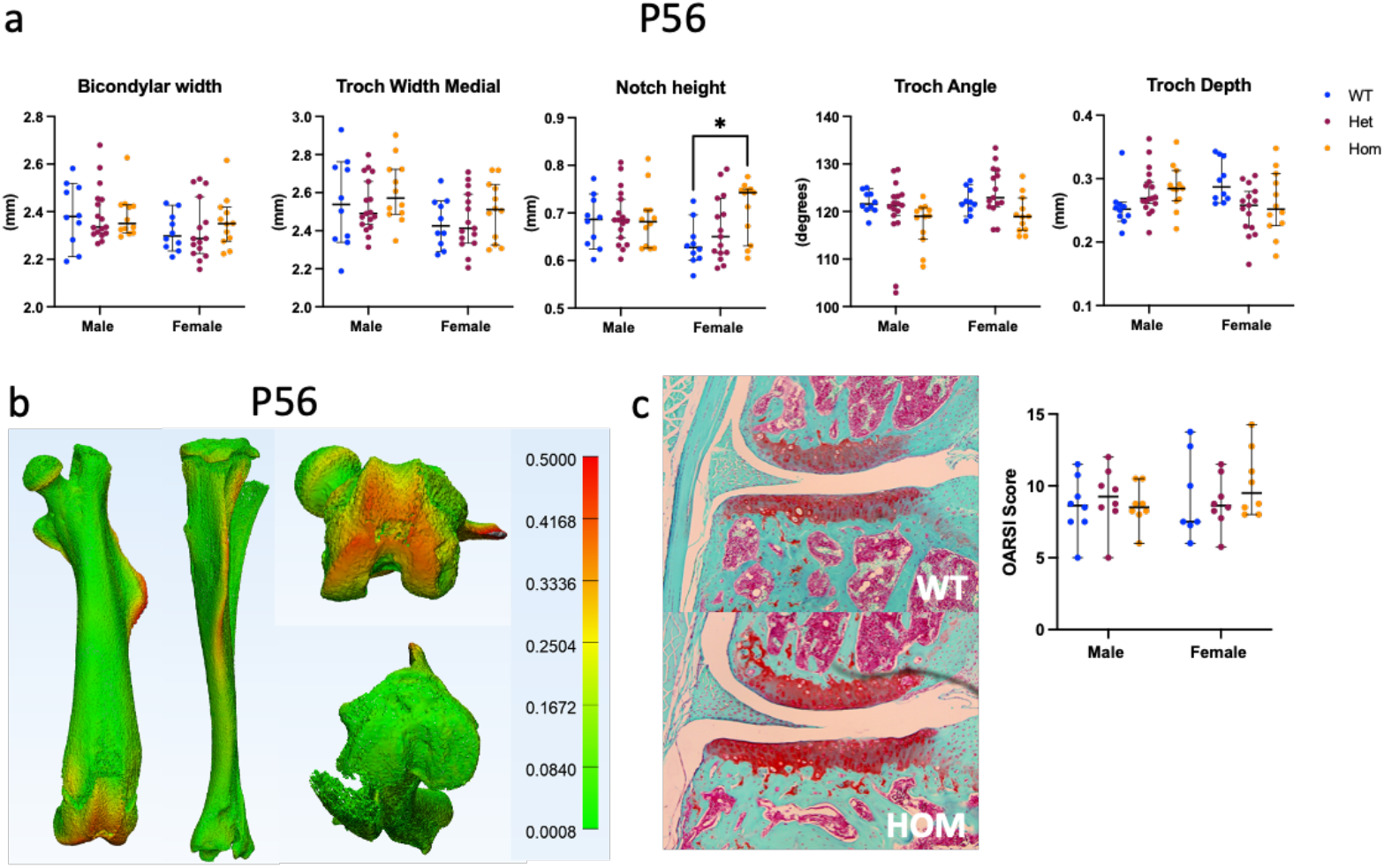
Morphological characterization of the *Gdf5 5’UTR* rs143384 variant in the mouse model. (a) MicroCT measurements of anatomical features in *5’UTR* rs143384 single base-pair replacement mice at P56 (Males: WT n = 10, Het n = 17, Hom n = 12. Females: WT n= 10, Het n= 15, Hom n= 12). Welch’s t test was used for comparisons between WT and Homozygous measurements (*p<0.05 bars indicate medians with 95% confidence intervals). (b) 3D morphological comparative analysis indicating locations of largest anatomical differences between WT and homozygous *risk 5’UTR* rs143384 hind limbs at P56. (c) Coronal histological sections of the medial compartment stained with Saf O of representative OARSI scoring (Males: WT n = 8, Het n = 8, Hom n = 8. Females: WT n= 7, Het n= 8, Hom n= 8).

### Modeling variant epistasis across the complex *GDF5* regulatory locus

The common 130 kb risk haplotype spanning *GDF5* associates with knee OA, DDH, height, and other musculoskeletal disease/traits (18). We had revealed causal risk alleles at rs4911178 (“A”) in *GROW1* and rs6060369 (“T”) in *R4*, uncoupling risk for DDH and knee OA, respectively. Yet above we revealed complex regulatory interactions within the vicinity of each variant and across the locus. Given that risk “A” rs143384 is in strong linkage disequilibrium with both risk “A” rs4911178 and “T” rs6060369 risk alleles, we hypothesized that there could be epistatic interactions between these variant positions (Fig. 1, variants in red). To first test for epistasis, we generated constructs containing different combinations of risk (R) and non-risk (N) alleles at three variants; rs143383 (*5’UTR*) (an additional associated variant), rs143384 (*5’UTR*), and rs6060369 (*R4*) (Fig. 8a) and transfected them into T/C-28a2 human chondrocytes (methods). We found that the *R4* risk/non-risk alleles drive lower luciferase expression than *5’UTR* risk/non-risk alleles. Strikingly, risk alleles at *R4* and *5’UTR* variants (Fig. 8a, blue values) drive nearly half the expression on non-risk alleles, indicating a strong epistatic interaction (Fig. 8a, red values). In the same fashion we tested for epistasis between or rs4911178 (in *GROW1*) and *5’UTR* variants, but transfected constructs into the human growth plate chondrocyte cell line, CHON-002. Here, we observe much lower luciferase expression with constructs containing either *GROW1* non-risk/risk alleles in comparison with *5’UTR* non-risk/risk alleles, with risk alleles driving lower expression than the non-risk. Additionally, we observe that all three risk alleles at *GROW1* and the *5’UTR* (Fig. 8b, red values) result again in an approximate halving of activity compared to non-risk alleles at these 3 variant positions (Fig. 8b, blue values). Together, these studies reveal complex interactions between variants that impact *GDF5* expression. We have included a diagram showing all interactions across the locus discovered to date (Fig. 8c).

**Figure 8.**
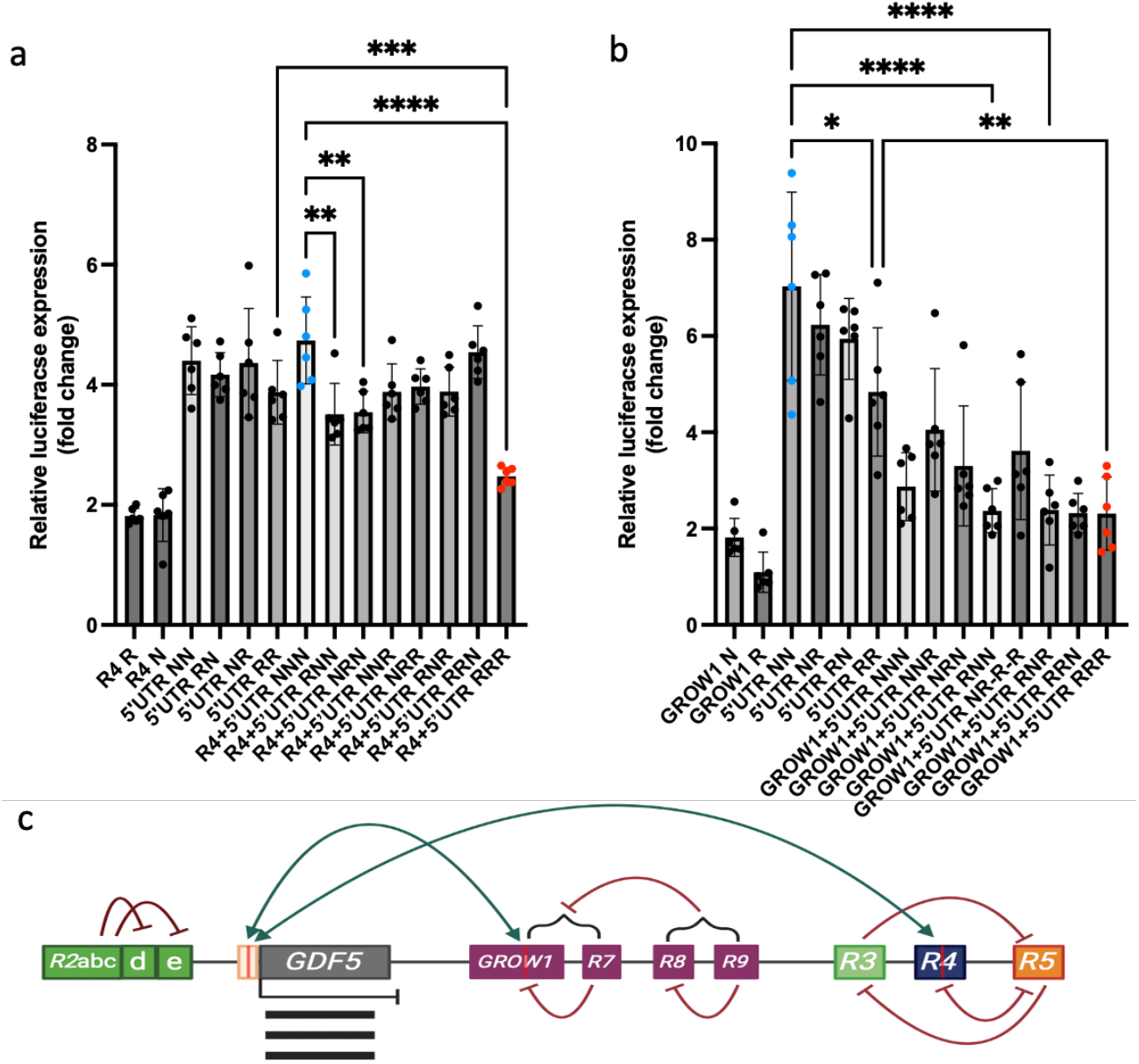
Disease risk variants interact to impact expression. (a) Relative normalized luciferase expression produced by plasmids containing different non-risk (N) or risk (R) variants of either the *R4* variant rs6060369, or the *5’UTR* variants rs143383 and rs143383 in T/C-28a2 cells. (b) Relative normalized luciferase expression produced by plasmids containing different non-risk (N) or risk (R) variants of either the *GROW1* variant rs4911178, or the *5’UTR* variants rs143383 and rs143383 in CHON-002 cells. (c) Cartoon model of identified interactions across the *GDF5* locus including risk variants marked as red lines (depictions not to scale).

## Discussion

### Localized interactions of regulatory sequences act to finely pattern Gdf5 expression

Prior to this study, what was known about *Gdf5* regulatory activity was restricted to broad patterns across the locus or specific regulatory sequences, with no large or small regulatory region able to fully recapitulate endogenous *Gdf5* expression. On interrogation of a downstream 37kb region encompassing the reported *GROW1* sub-perichondral enhancer (Fig.1, purple region), we identified an overlapping conserved subregion *PHC17* that drove *lacZ* expression throughout the growth plate, a pattern not observed for any larger or smaller construct nor for endogenous *Gdf5* expression. Indeed, *PHC17* individual components, *GROW1, R18-20*, and *R7*, were each unable to recapitulate this full growth plate signal. Interestingly, in the adjacent *PHC18* subregion, we found that the *R9* element acts to restrict *R8* expression patterns to only the shoulder joint, with its repressor activity confirmed in human chondrocytes. Therefore, *R9* is the strongest candidate to restrict *GROW1* to the sub-perichondral region across the entire 37 kb window. Further downstream in the 41kb joint region, we then observed that different combinations of the *R3, R4*, and *R5* enhancers regions drove very distinct patterns across limb joints with none alone able to recapitulate the complete 41kb, *PHC19*, or *Gdf5* expression patterns. Here, we identified that these enhancers also act as repressors with small or large impacts on expression depending on the skeletal site. Finally, for the upstream joint region *R2*, we identified subregion *abc* as a repressor sequence of the *d* and *e* subregions. Moreover, we identified a shorter 112bp sequence within the *d* region itself that restricts *d* expression to the major hind limb joints only. From these findings we conclude that there are complex activator and repressor sequences that operate between and within regulatory regions that can be tissue/cell type dependent as well as joint specific. We argue that this complex regulatory landscape likely evolved to precisely shape joints and long-bones, a point evinced by the high dysmorphic joints of *Gdf5* complete loss-of-function (*bp/bp)* mice.

### GDF5 complex regulatory landscape impacts how, when, and where risk variants drive abnormal disease phenotypes

By generating mice with a deletion of the *R2de* sequence adjacent to *Gdf5 5’UTR* we identified a 40% decrease per allele in *Gdf5* expression at all large limb joint skeletal sites. Interestingly, complete loss-of-function (*R2de*^*-/-*^ mice) did lead to hip and knee morphological changes early on (P56), but by 1.5 years these differences were no longer significant and importantly did not lead to any increase in OA disease. This is in-line with what is observed in mice with *Gdf5* coding mutations - i.e., *bp/+* have a 50% drop in expression and *bp/bp* mice have no functional expression, and neither develop OA, even though *bp/bp* mice have massive joint disruptions throughout life. These observations are in stark contrast to our findings from functional tests of individual *Gdf5* regulatory sequences and variants therein. Mice with a *R4* enhancer deletion have decreases (of ∼32%/allele) in *Gdf5* expression only in the knee, but not in other joint sites. This joint-specific decrease led to morphological changes in *R4*^*-/-*^ mice only in the knee at P56 and 1 year, with resulting effects on OA. Humanized mice with a GWAS OA risk variant (rs6060369, C→ T) in *R4* also have decreased *Gdf5* expression (∼16%/allele) within knee epiphyseal and articular cartilage only, and this reduction causes significant morphological changes in the same direction as OA patients, and a 30% increase in OA in homozygous “T/T” mice (9). Mice with a *GROW1* enhancer deletion have decreases (16%/allele) in *Gdf5* expression only in the proximal femur and acetabulum, but not at other joint sites. This led to femoral head and neck and acetabulum alterations (but not in knees), recapitulating those observed in DDH patients. Finally, humanized mice with a GWAS DDH risk variant (rs4911178, G→A) in *GROW1* also have decreased *Gdf5* expression (16%/allele) only within the proximal femur and acetabulum sub-perichondral chondrocytes, and this causes hip changes in mice in the same direction of effect as DDH patients (9).

Collectively, these functional studies reveal that location-specific decreases in *Gdf5* expression are far more important mediators of complex disease risk (OA, DDH, etc.) than large effects of expression reduction observed across different joint sites and tissues, the latter which are typically not observed in human patients with complex (i.e., common variant mediated) joint disease (but in fact are more indicative of patients with syndromic disorders due to *GDF5* coding mutations). We argue that this is the case because such location-specific decreases alter joint shape locally, changes that are not reciprocated on the opposing joint surface; resulting in a misregistration of joint articulation, which developmentally (DDH) or over time (OA), can predispose to disease. Furthermore, these effects on complex disease risk are in turn the product of the fine sculpting of *GDF5* expression by complex interacting regulatory sequences.

### Regulatory variant interactions drive complex disease risk at GDF5

We have shown that across the locus, GWAS risk variants can reside in enhancers (e.g., rs4911178 in *GROW1*), repressors (rs2378349 and rs2248393 in *R9*), or in sequences that behave as both (e.g., rs6060369 in *R4*), which in turn markedly obscures one’s ability to pinpoint causality at the individual base-pair level and at cell-type resolution. For example, by generating humanized mice harboring the most highly cited OA SNP variant (rs143384, G→A) located in *GDF5 5’UTR*, we did not observe any significant expression reductions at any joint site, nor did we find any significant morphological changes, or evidence of knee OA disease in adult mice. And this reveals that alone, rs143384 “A” is likely not causal for OA disease risk. Yet, we would be remiss to have not considered this (or any) variant within *GDF5* complex *cis*-regulatory architecture; a system that has evolved built-in activation/repression mechanisms locally (i.e., within a regulatory element) and further afield (across broad growth plate or joint regions) to generate endogenous *Gdf5* expression territories at each joint site, and thus precisely sculpt joints. Indeed, by testing for epistasis, we did observe substantial interactions between *5’UTR* rs143384 “A” and *R4* rs6060369 “T” risk alleles, and with *GROW1* rs4911178 “A” risk alleles. In each context, we argue that *cis*-regulatory variants provide locational and cell-type specificity, and by doing so they (i.e., *R4* and *GROW1*) both reduce and direct how reductions caused by other (*5’UTR*) variants become channeled to specific tissues. This in turn results in local joint mis-registration causing developmentally driven site-specific musculoskeletal disease risk at *GDF5*. We posit that disease risk at this developmental locus and most others, are analogous to the risks for developing cancers due to the actions of double or triple hits, but here hits are the myriad *cis*-regulatory variants residing on risk haplotypes and within their built-in complex epigenetic interactions (37, 38). Therefore, understanding the epistatic context in which a disease risk variant lies will be fundamental to disentangling and prioritizing the close to 2000 OA associated GWAS (39, 40) variants that have been identified.

## Acknowledgements

The authors would like to thank EpigenDx for ASE; Applied StemCell for mice; Dr. Li Zeng (Tufts University) and Dr. Mary Goldring (The Hospital for Special Surgery) for the T/C-28a2 cell line; Dr. Hopi Hoekstra for the NIH3T3 cell line; Dr. David Kingsley for advisement and guidance; and members of the Capellini Lab for support. This work was supported by NIH/NIAMS (1R01AR070139), Harvard University Dean’s Competitive Fund, Harvard University Milton Fund for Human Research to T.D.C; and Harvard University PRISE to S.Y; The Children’s Orthopaedic Surgery Foundation, Institutional Centers for Clinical and Translational Research at Boston Children’s Hospital, Harvard Clinical and Translational Science Center (National Center for Advancing Translational Sciences, NIH Award 1UM1TR004408-01), NIH (P30 AR075042) to A.K.

## Data Availability Statement

All data produced in the present work are contained in the manuscript.

