## Supplementary Materials for "Complex regulatory interactions at *GDF5* shape joint morphology and osteoarthritis disease risk"

**Supplementary Figures**

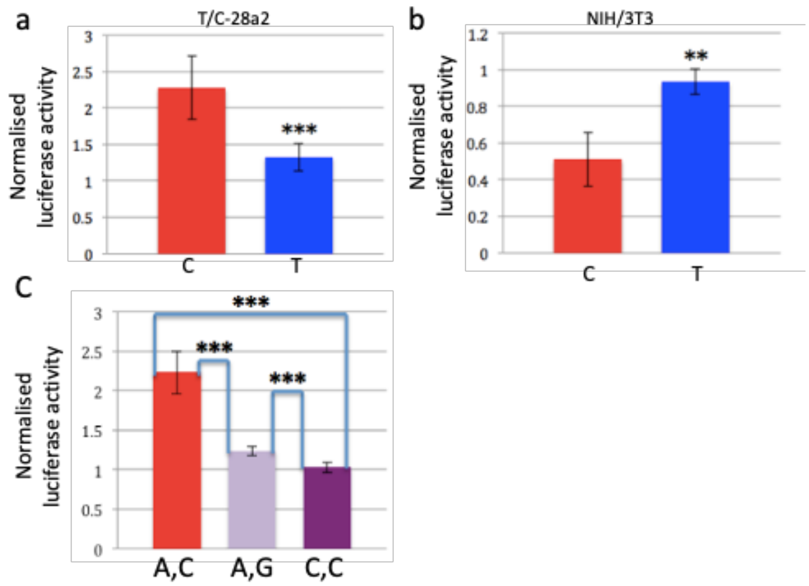


**Supplementary Figure 1.** Relative luciferase expression of plasmid constructs containing either the non-risk 'C' or risk 'T' allele of the *R4* rs6060369 risk position in (a) T/C-28a2 cells or (b) NIH/3T3 cells. (c) Relative luciferase expression of plasmid constructs targeting different combinations or risk and non-risk variants in *R9*; rs2378349 (non-risk "A", risk "C") and rs2248393 (risk "C", non-risk "G"), in comparison to empty vector (EV= 1). Error bars correspond to Mean ± SD, **p<0.001, ***p<0.0001. See Methods for replicate numbers.

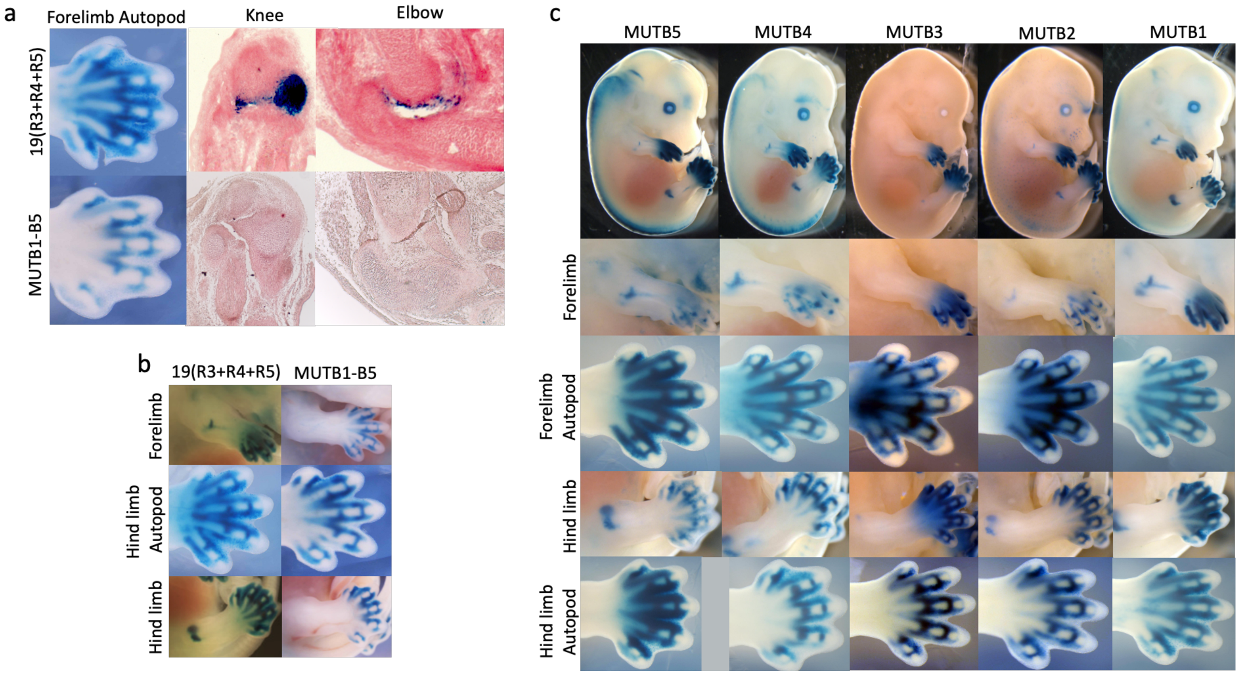


**Supplementary Figure 2.** (a) Transgenic E14.5 embryos with construct *PHC19* driving *lacZ* expression alongside construct *PHC19* containing 5 mutated BARX sites located within *R4* indicates loss of expression-driving capacity in the forelimb and hindlimb joints but not in the forelimb autopod. *LacZ* activity is shown in the forelimb autopod, and histological staining in the knee and elbow showing loss of activity where all 5 sites have undergone mutagenesis (MUTB1-5). (b) Further images of *lacZ* expression in the forelimb, hindlimb and hindlimb autopod of construct *PHC19* and constructs MUTB1-5. (c) Representative images of embryos with each independent BARX site mutated (MUTB5, MUTB4, MUTB3, MUTB2, MUTB1), with MUTB3 showing total loss of forelimb and hindlimb *lacZ* joint expression and MUTB2 having reduced expression. Mutagenesis of sites, MUTB5, MUTB4 and MUTB1 had no impact on *lacZ* expression in any joints.

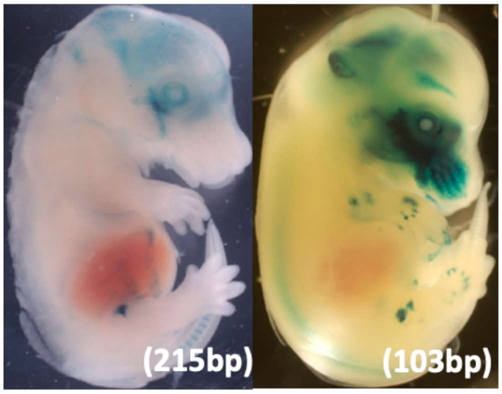


**Supplementary Figure 3.** Transgenic E14.5 *lacZ* embryos of subregion *d* (left) is 215bp in length. A portion of subregion *d* (right) that is 103bp in length drives expression in hindlimb and forelimb joints indicating that there are 112bp within subregion d that internally represses expression.

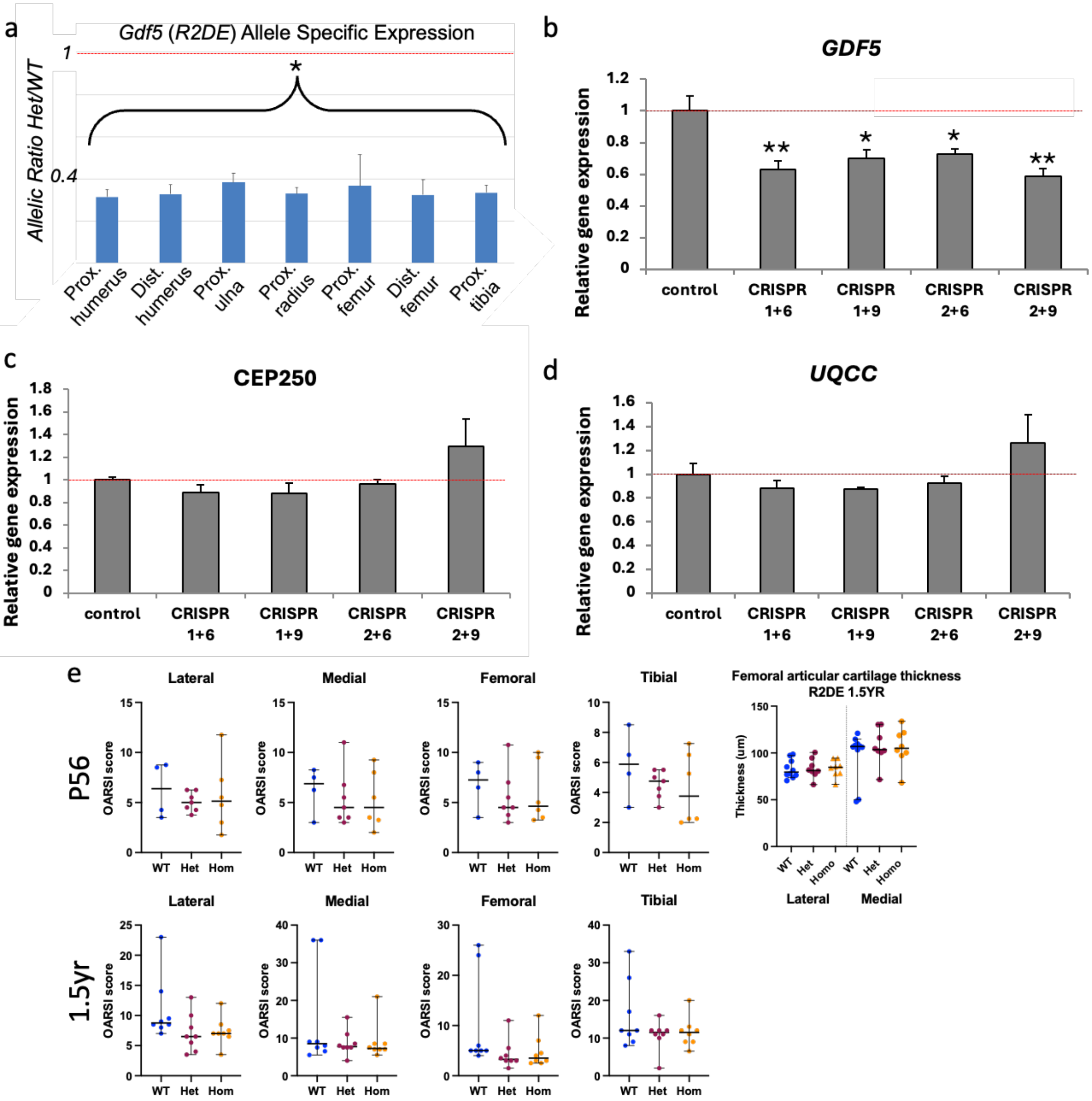
a
Gdf5 (R2DE) Allele Specific Expression
Allelic Ratio Het/WT
1
0.4
*
Prox. humerus
Dist. humerus
Prox. ulna
Prox. radius
Prox. femur
Dist. femur
Prox. tibia
b
GDF5
Relative gene expression
1.2
1
0.8
0.6
0.4
0.2
0
**
*
*
**
control
CRISPR 1+6
CRISPR 1+9
CRISPR 2+6
CRISPR 2+9
c
CEP250
Relative gene expression
1.8
1.6
1.4
1.2
1
0.8
0.6
0.4
0.2
0
control
CRISPR 1+6
CRISPR 1+9
CRISPR 2+6
CRISPR 2+9
d
UQCC
Relative gene expression
1.6
1.4
1.2
1
0.8
0.6
0.4
0.2
0
control
CRISPR 1+6
CRISPR 1+9
CRISPR 2+6
CRISPR 2+9
e
P56
Lateral
Medial
Femoral
Tibial
OARSI score
WT
Het
Hom
Femoral articular cartilage thickness R2DE 1.5YR
Thickness (um)
150
100
50
0
WT
Het
Homo
Lateral
Medial
1.5yr
Lateral
Medial
Femoral
Tibial
OARSI score
WT
Het
Hom

f

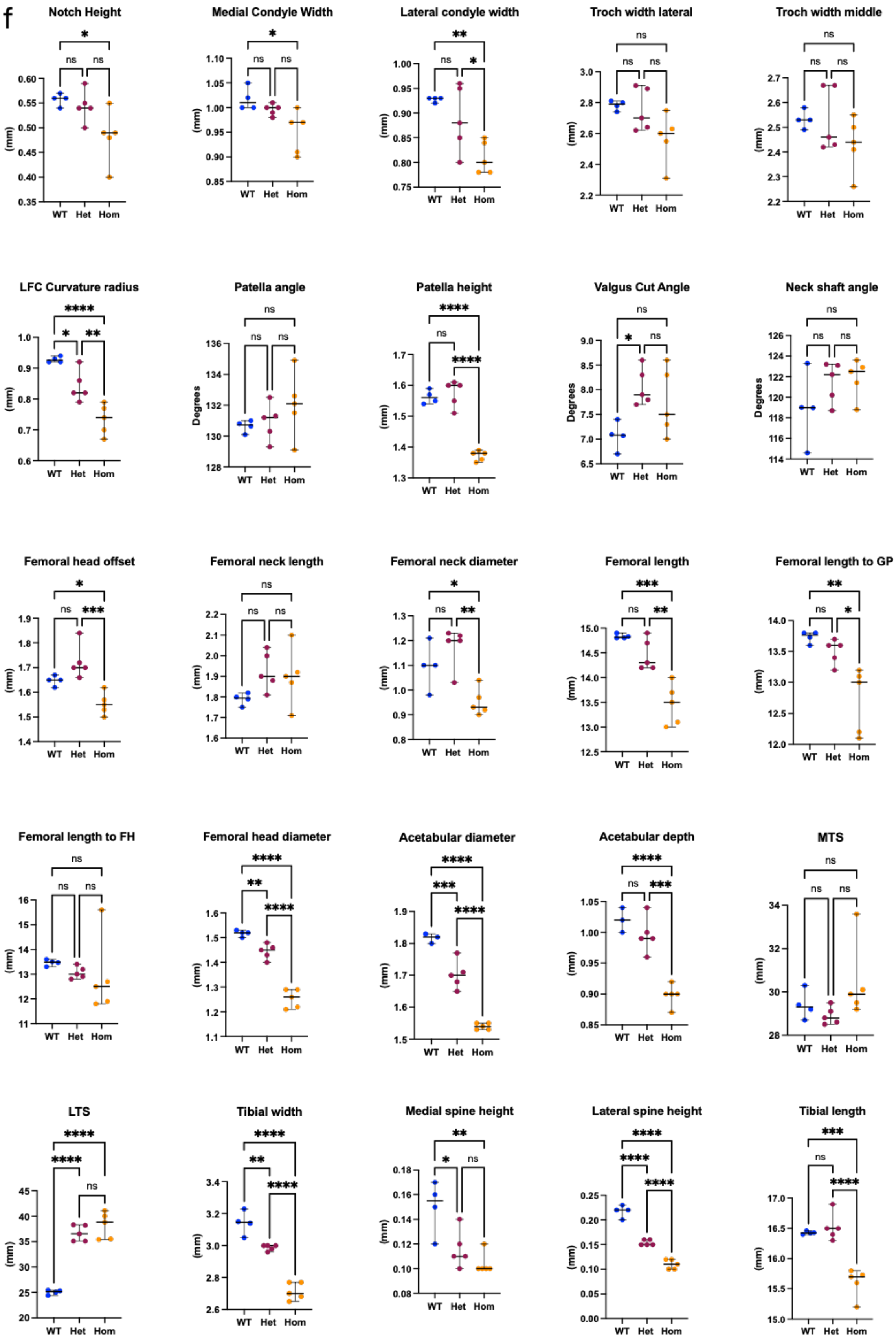

Notch Height
*
ns ns
0.60 0.55 0.50 0.45 0.40 0.35
(mm)
WT Het Hom
Medial Condyle Width
*
ns ns
1.10 1.05 1.00 0.95 0.90 0.85
(mm)
WT Het Hom
Lateral condyle width
**
ns *
1.00 0.95 0.90 0.85 0.80 0.75
(mm)
WT Het Hom
Troch width lateral
ns
ns ns
3.0 2.8 2.6 2.4 2.2
(mm)
WT Het Hom
Troch width middle
ns
ns ns
2.7 2.6 2.5 2.4 2.3 2.2
(mm)
WT Het Hom
LFC Curvature radius
****
* **
1.0 0.9 0.8 0.7 0.6
(mm)
WT Het Hom
Patella angle
ns
ns ns
136 134 132 130 128
Degrees
WT Het Hom
Patella height
****
ns ****
1.7 1.6 1.5 1.4 1.3
(mm)
WT Het Hom
Valgus Cut Angle
ns
* ns
9.0 8.5 8.0 7.5 7.0 6.5
Degrees
WT Het Hom
Neck shaft angle
ns
ns ns
126 124 122 120 118 116 114
Degrees
WT Het Hom
Femoral head offset
*
ns ***
1.9 1.8 1.7 1.6 1.5 1.4
(mm)
WT Het Hom
Femoral neck length
ns
ns ns
2.2 2.1 2.0 1.9 1.8 1.7 1.6
(mm)
WT Het Hom
Femoral neck diameter
*
ns **
1.3 1.2 1.1 1.0 0.9 0.8
(mm)
WT Het Hom
Femoral length
***
ns **
15.0 14.5 14.0 13.5 13.0 12.5
(mm)
WT Het Hom
Femoral length to GP
**
ns *
14.0 13.5 13.0 12.5 12.0
(mm)
WT Het Hom
Femoral length to FH
ns
ns ns
16 15 14 13 12 11
(mm)
WT Het Hom
Femoral head diameter
****
** ****
1.6 1.5 1.4 1.3 1.2 1.1
(mm)
WT Het Hom
Acetabular diameter
****
*** ****
1.9 1.8 1.7 1.6 1.5
(mm)
WT Het Hom
Acetabular depth
****
ns ***
1.05 1.00 0.95 0.90 0.85
(mm)
WT Het Hom
MTS
ns
ns ns
34 32 30 28
(mm)
WT Het Hom
LTS
****
**** ns
45 40 35 30 25 20
(mm)
WT Het Hom
Tibial width
****
** ****
3.4 3.2 3.0 2.8 2.6
(mm)
WT Het Hom
Medial spine height
**
* ns
0.18 0.16 0.14 0.12 0.10 0.08
(mm)
WT Het Hom
Lateral spine height
****
**** ****
0.25 0.20 0.15 0.10 0.05 0.00
(mm)
WT Het Hom
Tibial length
***
ns ****
17.0 16.5 16.0 15.5 15.0
(mm)
WT Het Hom

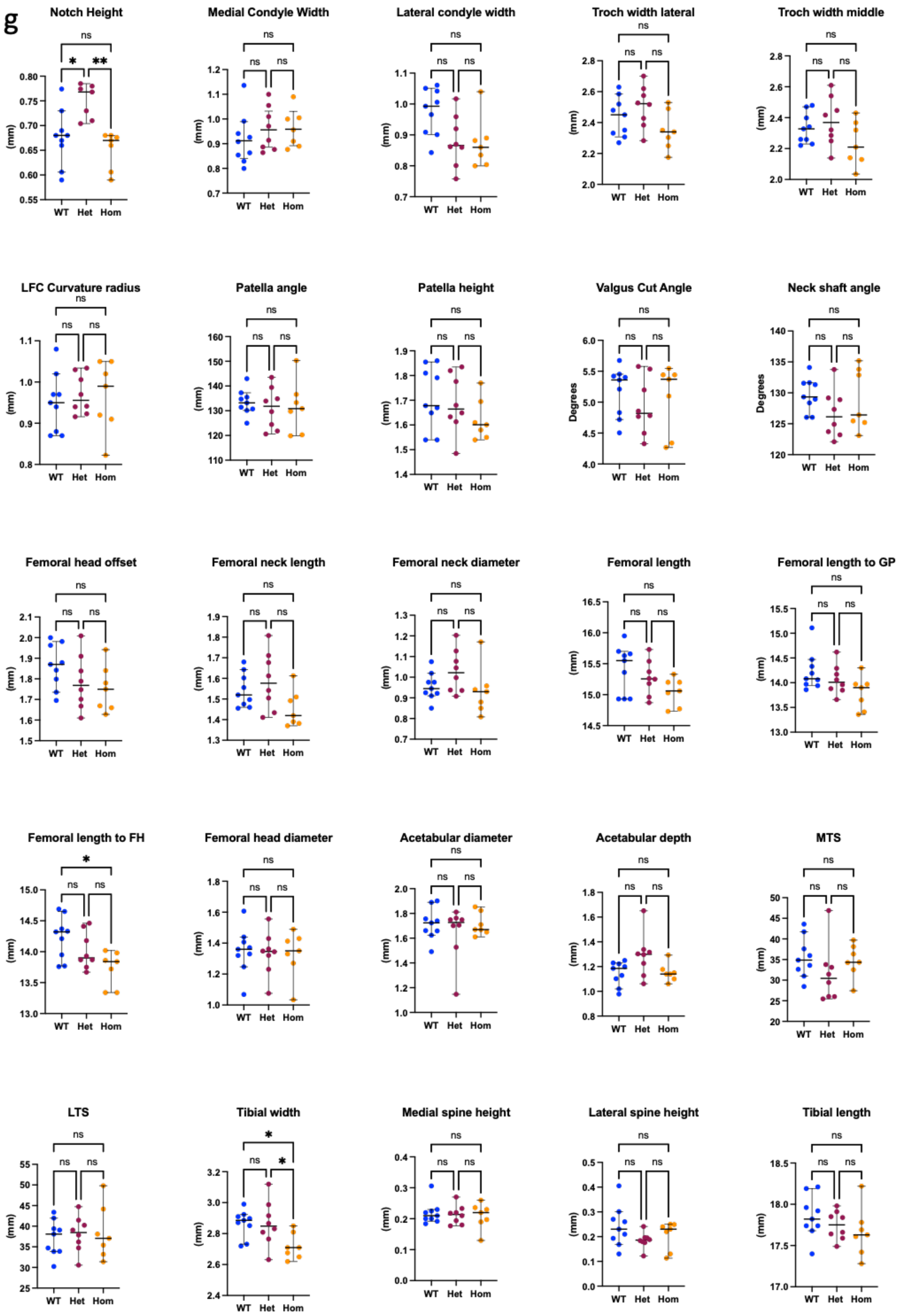
g
Notch Height
*
**
ns
(mm)
WT Het Hom
Medial Condyle Width
Lateral condyle width
Troch width lateral
Troch width middle
LFC Curvature radius
Patella angle
Patella height
Valgus Cut Angle
Degrees
Neck shaft angle
Femoral head offset
Femoral neck length
Femoral neck diameter
Femoral length
Femoral length to GP
Femoral length to FH
Femoral head diameter
Acetabular diameter
Acetabular depth
MTS
LTS
Tibial width
Medial spine height
Lateral spine height
Tibial length

**Supplementary Figure 4.** *In vitro* and *in vivo* targeting of the *R2de* regulatory region. (a) Allele-specific expression analysis at E15.5 reveals a 40% reduction in expression of *Gdf5* in all joint sites tested. Mean and standard deviation plotted. CRISPR guides targeting the *R2de* region in T/C-28a2 cells revealed reduction in expression of only (b) *GDF5* and not (c) upstream (*CEP250*) or (d) downstream (*UQCC*) genes within the *GDF5* risk locus; mean and standard deviation shown of qPCR replicate wells. (e) Femoral articular cartilage thickness measurements of the lateral and medial plateaus at P56 and 1.5 years alongside respective OARSI scores for each time point for the lateral, medial, femoral, and tibial plateaus (P56: WT n = 4, *R2de* Het = 5 *R2de* HOMO n = 6; 1.5 years: WT n= 8, Het n= 7, Hom n = 7). (f-g) Anatomical measurements of *R2de* wild-type (WT), Heterozygous (Het), and Homo (Homozygous) animals at P56 (f) and 1.5years (g). P56: (WT n = 4, Het n = 5, Hom n = 6) and 1.5 years: (WT n= 8, Het n= 7, Hom n = 7). *$p<0.05$, **$p<0.01$, ***$p<0.001$, ****$p<0.0001$, bars indicate medians and 95% Confidence Intervals.

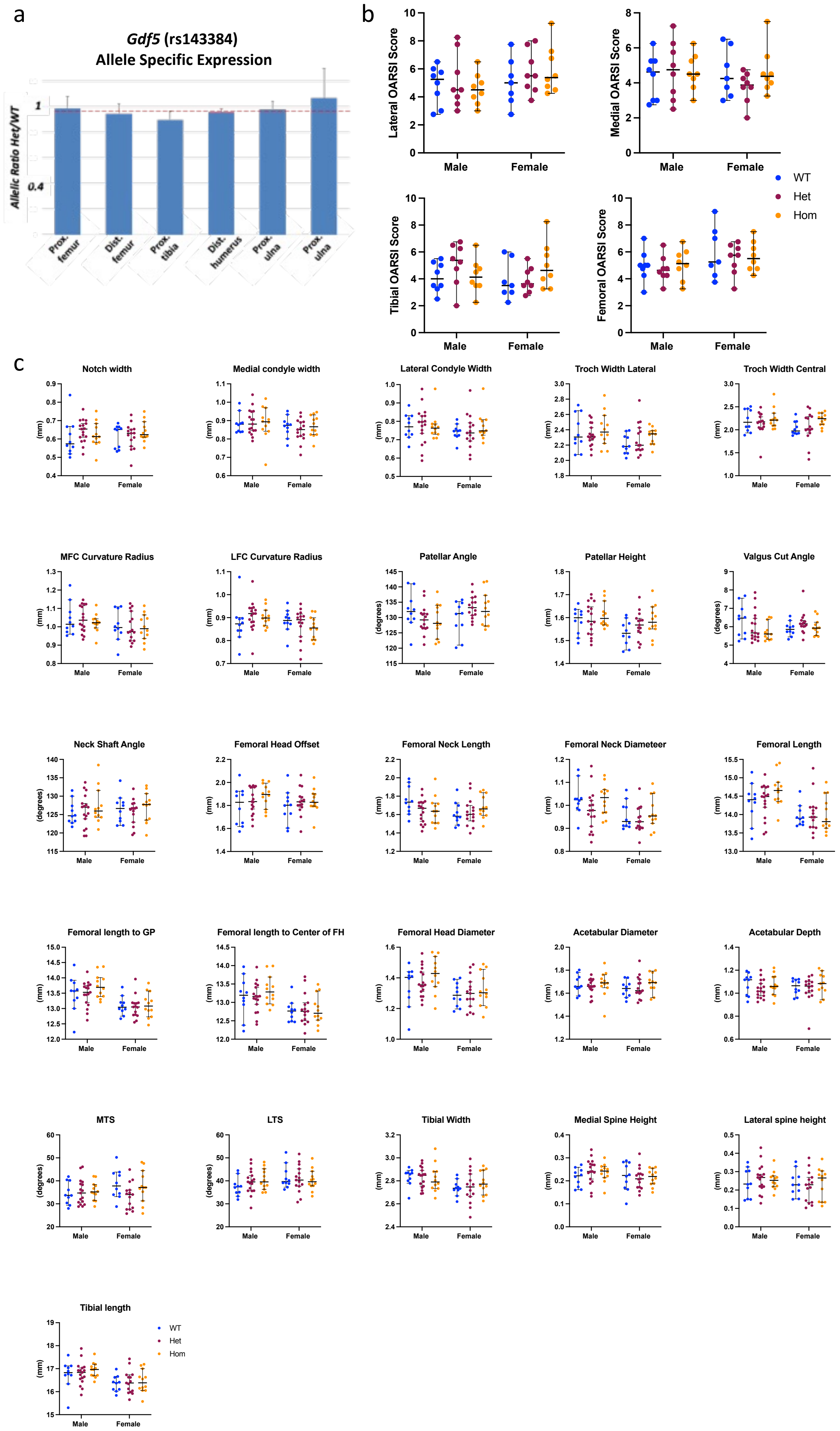
a
Gdf5 (rs143384)
Allele Specific Expression
Allelic Ratio Het/WT
1
0.4
Prox. femur
Dist. femur
Prox. tibia
Dist. humerus
Prox. ulna
Prox. ulna
b
Lateral OARSI Score
Medial OARSI Score
Tibial OARSI Score
Femoral OARSI Score
Male
Female
WT
Het
Hom
c
Notch width
Medial condyle width
Lateral Condyle Width
Troch Width Lateral
Troch Width Central
MFC Curvature Radius
LFC Curvature Radius
Patellar Angle
Patellar Height
Valgus Cut Angle
Neck Shaft Angle
Femoral Head Offset
Femoral Neck Length
Femoral Neck Diameteer
Femoral Length
Femoral length to GP
Femoral length to Center of FH
Femoral Head Diameter
Acetabular Diameter
Acetabular Depth
MTS
LTS
Tibial Width
Medial Spine Height
Lateral spine height
Tibial length
(mm)
(degrees)
Male
Female
WT
Het
Hom

**Supplementary Figure 5.** *In vivo* targeting of rs143384. (a) Allele-specific expression analysis at E15.5 revealed no effect of the 'T' risk allele on *Gdf5* expression in all joint sites tested; mean and standard deviation plotted. (b) Coronal histological OARSI scores of lateral, medial, tibial, and femoral plateaus at P56 reveal no regional specific effect of the *5'UTR* rs143384 risk variant (Males: WT n = 8, Het n = 8, Hom n = 8. Females: WT n= 7, Het n= 8, Hom n= 8). (c) Anatomical measurements of *5'UTR* rs143384 wild-type (WT), Heterozygous (Het) and Homozygous (Hom) animals in male and female mice at P56. Males: WT n = 10, Het n = 17, Hom n = 12. Females: WT n= 10, Het n= 15, Hom n= 12. *p<0.05, **p<0.01, ***p<0.001, ****p<0.0001, bars indicate medians and 95% Confidence Intervals.

### Supplementary Materials

### BARX binding sites

BARX sites (1-5) in *R4* regulatory element

TGTCCTCTACAGAGCATGTTGCCTATGAATACATAGCTCCCCCAGCAGAAG
ATGGCATGGGGCTGTTTGCTTATTAAAATCTTATCCTGTCCATAACGCTTTGC
TTTTTATAAAAACCCATACATACTGAGCCACCACATGGGAAGGGAGACCTT
TATATGTTACTCATATTTTCCCATTTATTTAAGCTATTAAAATGCATGCAGATA
TTTTATAAGCCATCAGGATTCGATGGCTGCTATTACTGAATGTTTAATAATAA
ACTAAACAGAAGAAATCAGAGGGTTAAATGATGACTGCAATTCCTGTAATG
ACTTAATTAAATTTTTATTGAGTACATAAACTATGATGTCACTTCTAGCCAAC
CGTGGCCCTGCTTTCTCAGCATGTCACAGGGTGGGAGCGCTCAATAGGAGG
CGAGGCTAAATGAGCAGCCGCTCTCTGCTGAGGTGGGGGAGGGAGCGAGG
CTCATGTAACACACTCAATAAATTATCCGTTCATTTTTAAACAAAGTCCATG
AGTTATAATATATTTGCCATGCACTTTTTATGACATAATAAAAATGTTACCTTA
AGGATTACTTAACACATACTACAAATGTTATAGCAGAGGCAGGCACAAGGG
GCAAAGAAAACAAACACGAGCGGGCCAGGGACGGGTCCTGGAGTTTATGT
CCCGGGGGAATGGCACAGTATATTAAAAGGGGAATGCAAAAAACAGGGGG
CACACGTGTTCAGGGTTACTAATCAGCAGATTAATGAATGAGACAGCCGAC
AGGGAGCACAACCAGAGGGGATCTTGCAAGGCACACATTTAATTAATGGA
AGTATTCTTAAGGAGGGACACGGGCTTTAAAGAGATGCCAGTTCATCTGAG
GCCAATCACTGGTCTGTCCCCTCTCACTGTTGCAGCCCTTAAGGACGTTTAT
AGACTGCTTAGCAGAGGATGAGTCCATGGTGTAGCCTGAAGGAGACT

N = BARX sites at E>.40

BARX sites (1-5) in *R4* regulatory element

TGTCCTCTACAGAGCATGTTGCCTATGAATACATAGCTCCCCCAGCAGAAG
ATGGCATGGGGCTGTTTGCTTATTAAAATCTTATCCTGTCCATAACGCTTTGC
TTTTTATAAAAACCCATACATACTGAGCCACCACATGGGAAGGGAGACCTT
TATATGTTACTCATATTTTCCCATTTCCGTACGCTCCGAACATGCATGCAGAT
ATTTTATAAGCCATCAGGATTCGATGGCTGCTATTACTGAATGTTTAATAATA
AACTAAACAGAAGAAATCAGAGGGTTAAATGATGACTGCAATTCCTGTAAT
GACTTACCGAACCGTTTATTGAGTACATAAACTATGATGTCACTTCTAGCCA
ACCGTGGCCCTGCTTTCTCAGCATGTCACAGGGTGGGAGCGCTCAATAGGA
GGCGAGGCTAAATGAGCAGCCGCTCTCTGCTGAGGTGGGGGAGGGAGCGA
GGCTCATGTAACACACTCAAGAACCGATCCGTTCATTTTTAAACAAAGTCC
ATGAGTTATAATATATTTGCCATGCACTTTTTATGACATAATAAAAATGTTACC
TTAAGGATTACTTAACACATACTACAAATGTTATAGCAGAGGCAGGCACAA
GGGGCAAAGAAAACAAACACGAGCGGGCCAGGGACGGGTCCTGGAGTTT
ATGTCCCGGGGGAATGGCACAGTATATTAAAAGGGGAATGCAAAAAACAG
GGGGCACACGTGTTCAGGGTTACTAATCAGCAGCCGAACGAATGAGACAG
CCGACAGGGAGCACAACCAGAGGGGATCTTGCAAGGCACACCCGTACCGA
ACGGAAGTATTCTTAAGGAGGGACACGGGCTTTAAAGAGATGCCAGTTCAT
CTGAGGCCAATCACTGGTCTGTCCCCTCTCACTGTTGCAGCCCTTAAGGAC
GTTTATAGACTGCTTAGCAGAGGATGAGTCCATGGTGTAGCCTGAAGGAGA
CT

N = mutated – no BARX sites at E>0.2

### Sequences of each risk and non-risk luciferase reporter construct

5'KpnI highlighted in Yellow. 3'HindIII highlighted in Cyan. NR= non-risk, R= risk.

1. GROW1_5UTR_CTT_NR-R-R

GGTACCTAAGGTACCTAAGCAGTAGTTTAGTAAGTGCACACCATATCCTGGGCACTTTGTGGGAACTAGGGAATAGTTCTCAAGAA
GCTCACAGTCTAGTGAGGGAGATAAGATGTGTAAAATGGTCAATTATAAATCAGAATGCCTAAGTTATATAATAAATATGAGTAAG
AAGTACTTATAGGATCACAAAGGAAGCTGCAACTATCAAAACCCTTATACTTCTACCTGTTAGTTCTTTGAAATGTGCCACGCATC
AGGTAGCTTAAAAGGCACACTGCCTGCCATCTTGTTGTCTGAATGGTGAAGGGTAAGGCATAGCGTTCAGGGAAGGAAGCTGCATA
TATTGAAGGGAAAAAGTGCTTTTTTAGGAGAGGAGGACTGTAGAGGTGGCACTGTAAAATTACTCACTGGAAAGGTGGGAGGGTTG
CAAGAAGGCTAGGGGAACTCAAAGATAATATTCAGAACCTGTATGAAAGACTCTTTATTGCCCATATAGCTAGAGGTATCAGTGTT
AGAGGGAGTGGATGGCAAAATAAAATAAAATATATAATCCCTGTGACCACATCCCAAGGTCAATACCTGACTAACAGCAGATGGGA
GAACCTTGAAACTACTTATAGTTCTTTAAAGTTTCCAAACTGTGCCTGCTTAATCTTCCTTGCTGAGGTTGGTTTAATTTCTGAGG
AGTAGGGTGTACCACCCCCAGTTCAATTGAGAGAGGGAAGCTGTCTGCTTCAAATCTATACTTGATTCCCCTTCCACTCTTTCTCT
CGCCTAGTTCTGATGATATGCAGGTTTTGGTGGAAAGAATCTGTAGCTTTTCAGTTTGAATCAGCACTCTTCTTGAGGACCTCCTA
TGTGTGCAGGACCATGGGATGTTGGAGAGAAGGGTATGACATGCATTCTTTTCTGGTAATTTCTTTTGGGCAAATCAGATACAGAC
TCTGTGTCTGTATCTTCAAGAAATGTAAGTCTAATGGAAAACAGACATGTTTAAACGGATTCAAAACTAGGGGGAAAAAAAAACTG
GAGCACACAGGCAGCATTACGCCATTCTTCCTTCTTGGAAAAATCCCTCAGCCTTATACAAGCCTCCTTCAAGCCCTCAGTCAGTT
GTGCAGGAGAAAGGGGGCGGTTGGCTTTCTCCTTTCAAGAACGAGTTATTTTCAGCTGCTGACTGGAGACGGTGCACGTCTGGATA
CGAGAGCATTTCCACTATGGGACTGGATACAAACACACACCCGGCAGACTTCAAGAGTCTCAGACTGAGGAGAAAGCCTTTCCTTC
TGCTGCTACTGCTGCTGCCGCTGCTTTTGAAAGTCCACTCCTTTCATGGTTTTTCCTGCCAAACCAGAGGCACCTTTGCTGCTGCC
GCTGTTCTCTTTGGTGTCATTCAGCGGAAGCTTGGCAAGCTT

2. GROW1_5UTR_CCT_NR-NR-R

GGTACCTAAGGTACCTAAGCAGTAGTTTAGTAAGTGCACACCATATCCTGGGCACTTTGTGGGAACTAGGGAATAGTTCTCAAGAA
GCTCACAGTCTAGTGAGGGAGATAAGATGTGTAAAATGGTCAATTATAAATCAGAATGCCTAAGTTATATAATAAATATGAGTAAG
AAGTACTTATAGGATCACAAAGGAAGCTGCAACTATCAAAACCCTTATACTTCTACCTGTTAGTTCTTTGAAATGTGCCACGCATC
AGGTAGCTTAAAAGGCACACTGCCTGCCATCTTGTTGTCTGAATGGTGAAGGGTAAGGCATAGCGTTCAGGGAAGGAAGCTGCATA
TATTGAAGGGAAAAAGTGCTTTTTTAGGAGAGGAGGACTGTAGAGGTGGCACTGTAAAATTACTCACTGGAAAGGTGGGAGGGTTG
CAAGAAGGCTAGGGGAACTCAAAGATAATATTCAGAACCTGTATGAAAGACTCTTTATTGCCCATATAGCTAGAGGTATCAGTGTT
AGAGGGAGTGGATGGCAAAATAAAATAAAATATATAATCCCTGTGACCACATCCCAAGGTCAATACCTGACTAACAGCAGATGGGA
GAACCTTGAAACTACTTATAGTTCTTTAAAGTTTCCAAACTGTGCCTGCTTAATCTTCCTTGCTGAGGTTGGTTTAATTTCTGAGG
AGTAGGGTGTACCACCCCCAGTTCAATTGAGAGAGGGAAGCTGTCTGCTTCAAATCTATACTTGATTCCCCTTCCACTCTTTCTCT

CGCCTAGTTCTGATGATATGCAGGTTTTGGTGGAAAGAATCTGTAGCTTTTCAGTTTGAATCAGCACTCTTCTTGAGGACCTCCTA
TGTGTGCAGGACCATGGGATGTTGGAGAGAAGGGTATGACATGCATTCTTTTCTGGTAATTTCTTTTGGGCAAATCAGATACAGAC
TCTGTGTCTGTATCTTCAAGAAATGTAAGTCTAATGGAAAACAGACATGTTTAAACGGATTCAAAACTAGGGGGAAAAAAAAACTG
GAGCACACAGGCAGCATTACGCCATTCTTCCTTCTTGGAAAAATCCCTCAGCCTTATACAAGCCTCCTTCAAGCCCTCAGTCAGTT
GTGCAGGAGAAAGGGGGCGGTCGGCTTTCTCCTTTCAAGAACGAGTTATTTTCAGCTGCTGACTGGAGACGGTGCACGTCTGGATA
CGAGAGCATTTCCACTATGGGACTGGATACAAACACACACCCGGCAGACTTCAAGAGTCTCAGACTGAGGAGAAAGCCTTTCCTTC
TGCTGCTACTGCTGCTGCCGCTGCTTTTGAAAGTCCACTCCTTTCATGGTTTTTCCTGCCAAACCAGAGGCACCTTTGCTGCTGCC
GCTGTTCTCTTTGGTGTCATTCAGCGGAAGCTTGGCAAGCTT

3. GROW1_5UTR_CTC_NR-R-NR
GGTACCTAAGGTACCTAAGCAGTAGTTTAGTAAGTGCACACCATATCCTGGGCACTTTGTGGGAACTAGGGAATAGTTCTCAAGAA
GCTCACAGTCTAGTGAGGGAGATAAGATGTGTAAAATGGTCAATTATAAATCAGAATGCCTAAGTTATATAATAAATATGAGTAAG
AAGTACTTATAGGATCACAAAGGAAGCTGCAACTATCAAAACCCCTTATACTTCTACCTGTTAGTTCTTTGAAATGTGCCACGCATC
AGGTAGCTTAAAAGGCACACTGCCTGCCATCTTGTTGTCTGAATGGTGAAGGGTAAGGCATAGCGTTCAGGGAAGGAAGCTGCATA
TATTGAAGGGAAAAAGTGCTTTTTTAGGAGAGGAGGACTGTAGAGGTGGCACTGTAAAATTACTCACTGGAAAGGTGGGAGGGTTG
CAAGAAGGCTAGGGGAACTCAAAGATAATATTCAGAACCTGTATGAAAGACTCTTTATTGCCCATATAGCTAGAGGTATCAGTGTT
AGAGGGAGTGGATGGCAAAATAAAATAAAATATATAATCCCTGTGACCACATCCCAAGGTCAATACCTGACTAACAGCAGATGGGA
GAACCTTGAAACTACTTATAGTTCTTTAAAGTTTCCAAACTGTGCCTGCTTAATCTTCCTTGCTGAGGTTGGTTTAATTTCTGAGG
AGTAGGGTGTACCACCCCCAGTTCAATTGAGAGAGGGAAGCTGTCTGCTTCAAATCTATACTTGATTCCCCTTCCACTCTTTCTCC
GCCTAGTTCTGATGATATGCAGGTTTTGGTGGAAAGAATCTGTAGCTTTTCAGTTTGAATCAGCACTCTTCTTGAGGACCTCCTAT
GTGTGCAGGACCATGGGATGTTGGAGAGAAGGGTATGACATGCATTCTTTTCTGGTAATTTCTTTTGGGCAAATCAGATACAGACT
CTGTGTCTGTATCTTCAAGAAATGTAAGTCTAATGGAAAACAGACATGTTTAAACGGATTCAAAACTAGGGGGAAAAAAAAACTGG
AGCACACAGGCAGCATTACGCCATTCTTCCTTCTTGGAAAAATCCCTCAGCCTTATACAAGCCTCCTTCAAGCCCTCAGTCAGTTG
TGCAGGAGAAAGGGGGCGGTTGGCTTTCTCCTTTCAAGAACGAGTTATTTTCAGCTGCTGACTGGAGACGGTGCACGTCTGGATAC
GAGAGCATTTCCACTATGGGACTGGATACAAACACACACCCGGCAGACTTCAAGAGTCTCAGACTGAGGAGAAAGCCTTTCCTTCT
GCTGCTACTGCTGCTGCCGCTGCTTTTGAAAGTCCACTCCTTTCATGGTTTTTCCTGCCAAACCAGAGGCACCTTCGCTGCTGCCG
CTGTTCTCTTTGGTGTCATTCAGCGGAAGCTTGGCAAGCTT

4. GROW1_5UTR_CCC_NR-NR-NR
GGTACCTAAGGTACCTAAGCAGTAGTTTAGTAAGTGCACACCATATCCTGGGCACTTTGTGGGAACTAGGGAATAGTTCTCAAGAA
GCTCACAGTCTAGTGAGGGAGATAAGATGTGTAAAATGGTCAATTATAAATCAGAATGCCTAAGTTATATAATAAATATGAGTAAG
AAGTACTTATAGGATCACAAAGGAAGCTGCAACTATCAAAACCCCTTATACTTCTACCTGTTAGTTCTTTGAAATGTGCCACGCATC
AGGTAGCTTAAAAGGCACACTGCCTGCCATCTTGTTGTCTGAATGGTGAAGGGTAAGGCATAGCGTTCAGGGAAGGAAGCTGCATA
TATTGAAGGGAAAAAGTGCTTTTTTAGGAGAGGAGGACTGTAGAGGTGGCACTGTAAAATTACTCACTGGAAAGGTGGGAGGGTTG
CAAGAAGGCTAGGGGAACTCAAAGATAATATTCAGAACCTGTATGAAAGACTCTTTATTGCCCATATAGCTAGAGGTATCAGTGTT
AGAGGGAGTGGATGGCAAAATAAAATAAAATATATAATCCCTGTGACCACATCCCAAGGTCAATACCTGACTAACAGCAGATGGGA
GAACCTTGAAACTACTTATAGTTCTTTAAAGTTTCCAAACTGTGCCTGCTTAATCTTCCTTGCTGAGGTTGGTTTAATTTCTGAGG
AGTAGGGTGTACCACCCCCAGTTCAATTGAGAGAGGGAAGCTGTCTGCTTCAAATCTATACTTGATTCCCCTTCCACTCTTTCTCT
CGCCTAGTTCTGATGATATGCAGGTTTTGGTGGAAAGAATCTGTAGCTTTTCAGTTTGAATCAGCACTCTTCTTGAGGACCTCCTA
TGTGTGCAGGACCATGGGATGTTGGAGAGAAGGGTATGACATGCATTCTTTTCTGGTAATTTCTTTTGGGCAAATCAGATACAGAC
TCTGTGTCTGTATCTTCAAGAAATGTAAGTCTAATGGAAAACAGACATGTTTAAACGGATTCAAAACTAGGGGGAAAAAAAAACTG
GAGCACACAGGCAGCATTACGCCATTCTTCCTTCTTGGAAAAATCCCTCAGCCTTATACAAGCCTCCTTCAAGCCCTCAGTCAGTT
GTGCAGGAGAAAGGGGGCGGTCGGCTTTCTCCTTTCAAGAACGAGTTATTTTCAGCTGCTGACTGGAGACGGTGCACGTCTGGATA
CGAGAGCATTTCCACTATGGGACTGGATACAAACACACACCCGGCAGACTTCAAGAGTCTCAGACTGAGGAGAAAGCCTTTCCTTC
TGCTGCTACTGCTGCTGCCGCTGCTTTTGAAAGTCCACTCCTTTCATGGTTTTTCCTGCCAAACCAGAGGCACCTTCGCTGCTGCC
GCTGTTCTCTTTGGTGTCATTCAGCGGAAGCTTGGCAAGCTT

5. GROW1_5UTR_TTT_R-R-R
GGTACCTAAGGTACCTAAGCAGTAGTTTAGTAAGTGCACACCATATCCTGGGCACTTTGTGGGAACTAGGGAATAGTTCTCAAGAA
GCTCACAGTCTAGTGAGGGAGATAAGATGTGTAAAATGGTCAATTATAAATCAGAATGCCTAAGTTATATAATAAATATGAGTAAG
AAGTACTTATAGGATCACAAAGGAAGCTGCAACTATCAAAACCCCTTATACTTCTACCTGTTAGTTCTTTGAAATGTGCCACGCATC
AGGTAGCTTAAAAGGCACACTGCCTGCCATCTTGTTGTCTGAATGGTGAAGGGTAAGGCATAGCGTTCAGGGAAGGAAGCTGCATA
TATTGAAGGGAAAAAGTGCTTTTTTAGGAGAGGAGGACTGTAGAGGTGGCACTGTAAAATTACTCACTGGAAAGGTGGGAGGGTTG
CAAGAAGGCTAGGGGAACTCAAAGATAATATTCAGAACCTGTATGAAAGACTCTTTATTGCCCATATAGCTAGAGGTATCAGTGTT
AGAGGGAGTGGATGGCAAAATAAAATAAAATATATAATCTCTGTGACCACATCCCAAGGTCAATACCTGACTAACAGCAGATGGGA
GAACCTTGAAACTACTTATAGTTCTTTAAAGTTTCCAAACTGTGCCTGCTTAATCTTCCTTGCTGAGGTTGGTTTAATTTCTGAGG
AGTAGGGTGTACCACCCCCAGTTCAATTGAGAGAGGGAAGCTGTCTGCTTCAAATCTATACTTGATTCCCCTTCCACTCTTTCTCT
CGCCTAGTTCTGATGATATGCAGGTTTTGGTGGAAAGAATCTGTAGCTTTTCAGTTTGAATCAGCACTCTTCTTGAGGACCTCCTA
TGTGTGCAGGACCATGGGATGTTGGAGAGAAGGGTATGACATGCATTCTTTTCTGGTAATTTCTTTTGGGCAAATCAGATACAGAC
TCTGTGTCTGTATCTTCAAGAAATGTAAGTCTAATGGAAAACAGACATGTTTAAACGGATTCAAAACTAGGGGGAAAAAAAAACTG
GAGCACACAGGCAGCATTACGCCATTCTTCCTTCTTGGAAAAATCCCTCAGCCTTATACAAGCCTCCTTCAAGCCCTCAGTCAGTT
GTGCAGGAGAAAGGGGGCGGTTGGCTTTCTCCTTTCAAGAACGAGTTATTTTCAGCTGCTGACTGGAGACGGTGCACGTCTGGATA

CGAGAGCATTTCCACTATGGGACTGGATACAAACACACACCCGGCAGACTTCAAGAGTCTCAGACTGAGGAGAAAGCCTTTCCTTC
TGCTGCTACTGCTGCTGCCGCTGCTTTTGAAAGTCCACTCCTTTCATGGTTTTTCCTGCCAAACCAGAGGCACCTTTGCTGCTGCC
GCTGTTCTCTTTGGTGTCATTCAGCGGAAGCTTGGCAAGCTT

6. GROW1_5UTR_TCT_R-NR-R
GGTACCTAAGGTACCTAAGCAGTAGTTTAGTAAGTGCACACCATATCCTGGGCACTTTGTGGGAACTAGGGAATAGTTCTCAAGAA
GCTCACAGTCTAGTGAGGGAGATAAGATGTGTAAAATGGTCAATTATAAATCAGAATGCCTAAGTTATATAATAAATATGAGTAAG
AAGTACTTATAGGATCACAAAGGAAGCTGCAACTATCAAAACCCTTATACTTCTACCTGTTAGTTCTTTGAAATGTGCCACGCATC
AGGTAGCTTAAAAGGCACACTGCCTGCCATCTTGTTGTCTGAATGGTGAAGGGTAAGGCATAGCGTTCAGGGAAGGAAGCTGCATA
TATTGAAGGGAAAAAGTGCTTTTTTAGGAGAGGAGGACTGTAGAGGTGGCACTGTAAAATTACTCACTGGAAAGGTGGGAGGGTTG
CAAGAAGGCTAGGGGAACTCAAAGATAATATTCAGAACCTGTATGAAAGACTCTTTATTGCCCATATAGCTAGAGGTATCAGTGTT
AGAGGGAGTGGATGGCAAAATAAAATAAAATATATAATCTCTGTGACCACATCCCAAGGTCAATACCTGACTAACAGCAGATGGGA
GAACCTTGAAACTACTTATAGTTCTTTAAAGTTTCCAAACTGTGCCTGCTTAATCTTCCTTGCTGAGGTTGGTTTAATTTCTGAGG
AGTAGGGTGTACCACCCCCAGTTCAATTGAGAGAGGGAAGCTGTCTGCTTCAAATCTATACTTGATTCCCCTTCCACTCTTTCTCT
CGCCTAGTTCTGATGATATGCAGGTTTTGGTGGAAAGAATCTGTAGCTTTTCAGTTTGAATCAGCACTCTTCTTGAGGACCTCCTA
TGTGTGCAGGACCATGGGATGTTGGAGAGAAGGGTATGACATGCATTCTTTTCTGGTAATTTCTTTTGGGCAAATCAGATACAGAC
TCTGTGTCTGTATCTTCAAGAAATGTAAGTCTAATGGAAAACAGACATGTTTAAACGGATTCAAAACTAGGGGGAAAAAAAAACTG
GAGCACACAGGCAGCATTACGCCATTCTTCCTTCTTGGAAAAATCCCTCAGCCTTATACAAGCCTCCTTCAAGCCCTCAGTCAGTT
GTGCAGGAGAAAGGGGGCGGTCGGCTTTCTCCTTTCAAGAACGAGTTATTTTCAGCTGCTGACTGGAGACGGTGCACGTCTGGATA
CGAGAGCATTTCCACTATGGGACTGGATACAAACACACACCCGGCAGACTTCAAGAGTCTCAGACTGAGGAGAAAGCCTTTCCTTC
TGCTGCTACTGCTGCTGCCGCTGCTTTTGAAAGTCCACTCCTTTCATGGTTTTTCCTGCCAAACCAGA
GGCACCTTTGCTGCTGCCGCTGTTCTCTTTGGTGTCATTCAGCGGAAGCTTGGCAAGCTT

7. GROW1_5UTR_TTC_R-R-NR
GGTACCTAAGGTACCTAAGCAGTAGTTTAGTAAGTGCACACCATATCCTGGGCACTTTGTGGGAACTAGGGAATAGTTCTCAAGAA
GCTCACAGTCTAGTGAGGGAGATAAGATGTGTAAAATGGTCAATTATAAATCAGAATGCCTAAGTTATATAATAAATATGAGTAAG
AAGTACTTATAGGATCACAAAGGAAGCTGCAACTATCAAAACCCTTATACTTCTACCTGTTAGTTCTTTGAAATGTGCCACGCATC
AGGTAGCTTAAAAGGCACACTGCCTGCCATCTTGTTGTCTGAATGGTGAAGGGTAAGGCATAGCGTTCAGGGAAGGAAGCTGCATA
TATTGAAGGGAAAAAGTGCTTTTTTAGGAGAGGAGGACTGTAGAGGTGGCACTGTAAAATTACTCACTGGAAAGGTGGGAGGGTTG
CAAGAAGGCTAGGGGAACTCAAAGATAATATTCAGAACCTGTATGAAAGACTCTTTATTGCCCATATAGCTAGAGGTATCAGTGTT
AGAGGGAGTGGATGGCAAAATAAAATAAAATATATAATCTCTGTGACCACATCCCAAGGTCAATACCTGACTAACAGCAGATGGGA
GAACCTTGAAACTACTTATAGTTCTTTAAAGTTTCCAAACTGTGCCTGCTTAATCTTCCTTGCTGAGGTTGGTTTAATTTCTGAGG
AGTAGGGTGTACCACCCCCAGTTCAATTGAGAGAGGGAAGCTGTCTGCTTCAAATCTATACTTGATTCCCCTTCCACTCTTTCTCT
CGCCTAGTTCTGATGATATGCAGGTTTTGGTGGAAAGAATCTGTAGCTTTTCAGTTTGAATCAGCACTCTTCTTGAGGACCTCCTA
TGTGTGCAGGACCATGGGATGTTGGAGAGAAGGGTATGACATGCATTCTTTTCTGGTAATTTCTTTTGGGCAAATCAGATACAGAC
TCTGTGTCTGTATCTTCAAGAAATGTAAGTCTAATGGAAAACAGACATGTTTAAACGGATTCAAAACTAGGGGGAAAAAAAAACTG
GAGCACACAGGCAGCATTACGCCATTCTTCCTTCTTGGAAAAATCCCTCAGCCTTATACAAGCCTCCTTCAAGCCCTCAGTCAGTT
GTGCAGGAGAAAGGGGGCGGTTGGCTTTCTCCTTTCAAGAACGAGTTATTTTCAGCTGCTGACTGGAGACGGTGCACGTCTGGATA
CGAGAGCATTTCCACTATGGGACTGGATACAAACACACACCCGGCAGACTTCAAGAGTCTCAGACTGAGGAGAAAGCCTTTCCTTC
TGCTGCTACTGCTGCTGCCGCTGCTTTTGAAAGTCCACTCCTTTCATGGTTTTTCCTGCCAAACCAGAGGCACCTTCGCTGCTGCC
GCTGTTCTCTTTGGTGTCATTCAGCGGAAGCTTGGCAAGCTT

8. GROW1_5UTR_TCC_R-NR-NR
GGTACCTAAGGTACCTAAGCAGTAGTTTAGTAAGTGCACACCATATCCTGGGCACTTTGTGGGAACTAGGGAATAGTTCTCAAGAA
GCTCACAGTCTAGTGAGGGAGATAAGATGTGTAAAATGGTCAATTATAAATCAGAATGCCTAAGTTATATAATAAATATGAGTAAG
AAGTACTTATAGGATCACAAAGGAAGCTGCAACTATCAAAACCCTTATACTTCTACCTGTTAGTTCTTTGAAATGTGCCACGCATC
AGGTAGCTTAAAAGGCACACTGCCTGCCATCTTGTTGTCTGAATGGTGAAGGGTAAGGCATAGCGTTCAGGGAAGGAAGCTGCATA
TATTGAAGGGAAAAAGTGCTTTTTTAGGAGAGGAGGACTGTAGAGGTGGCACTGTAAAATTACTCACTGGAAAGGTGGGAGGGTTG
CAAGAAGGCTAGGGGAACTCAAAGATAATATTCAGAACCTGTATGAAAGACTCTTTATTGCCCATATAGCTAGAGGTATCAGTGTT
AGAGGGAGTGGATGGCAAAATAAAATAAAATATATAATCTCTGTGACCACATCCCAAGGTCAATACCTGACTAACAGCAGATGGGA
GAACCTTGAAACTACTTATAGTTCTTTAAAGTTTCCAAACTGTGCCTGCTTAATCTTCCTTGCTGAGGTTGGTTTAATTTCTGAGG
AGTAGGGTGTACCACCCCCAGTTCAATTGAGAGAGGGAAGCTGTCTGCTTCAAATCTATACTTGATTCCCCTTCCACTCTTTCTCT
CGCCTAGTTCTGATGATATGCAGGTTTTGGTGGAAAGAATCTGTAGCTTTTCAGTTTGAATCAGCACTCTTCTTGAGGACCTCCTA
TGTGTGCAGGACCATGGGATGTTGGAGAGAAGGGTATGACATGCATTCTTTTCTGGTAATTTCTTTTGGGCAAATCAGATACAGAC
TCTGTGTCTGTATCTTCAAGAAATGTAAGTCTAATGGAAAACAGACATGTTTAAACGGATTCAAAACTAGGGGGAAAAAAAAACTG
GAGCACACAGGCAGCATTACGCCATTCTTCCTTCTTGGAAAAATCCCTCAGCCTTATACAAGCCTCCTTCAAGCCCTCAGTCAGTT
GTGCAGGAGAAAGGGGGCGGTCGGCTTTCTCCTTTCAAGAACGAGTTATTTTCAGCTGCTGACTGGAGACGGTGCACGTCTGGATA
CGAGAGCATTTCCACTATGGGACTGGATACAAACACACACCCGGCAGACTTCAAGAGTCTCAGACTGAGGAGAAAGCCTTTCCTTC
TGCTGCTACTGCTGCTGCCGCTGCTTTTGAAAGTCCACTCCTTTCATGGTTTTTCCTGCCAAACCAGAGGCACCTTCGCTGCTGCC
GCTGTTCTCTTTGGTGTCATTCAGCGGAAGCTTGGCAAGCTT

9. GROW1_NR

GGTACCTAAGGTACCTAAGCAGTAGTTTAGTAAGTGCACACCATATCCTGGGCACTTTGTGGGAACTAGGGAATAGTTCTCAAGAA
GCTCACAGTCTAGTGAGGGAGATAAGATGTGTAAAATGGTCAATTATAAATCAGAATGCCTAAGTTATATAATAAATATGAGTAAG
AAGTACTTATAGGATCACAAAGGAAGCTGCAACTATCAAAACCCTTATACTTCTACCTGTTAGTTCTTTGAAATGTGCCACGCATC
AGGTAGCTTAAAAGGCACACTGCCTGCCATCTTGTTGTCTGAATGGTGAAGGGTAAGGCATAGCGTTCAGGGAAGGAAGCTGCATA
TATTGAAGGGAAAAAGTGCTTTTTTAGGAGAGGAGGACTGTAGAGGTGGCACTGTAAAATTACTCACTGGAAAGGTGGGAGGGTTG
CAAGAAGGCTAGGGGAACTCAAAGATAATATTCAGAACCTGTATGAAAGACTCTTTATTGCCCATATAGCTAGAGGTATCAGTGTT
AGAGGGAGTGGATGGCAAAATAAAATAAAATATATAATCCCTGTGACCACATCCCAAGGTCAATACCTGACTAACAGCAGATGGGA
GAACCTTGAAACTACTTATAGTTCTTTAAAGTTTCCAAACTGTGCCTGCTTAATCTTCCTTGCTGAGGTTGGTTTAATTTCTGAGG
AGTAGGGTGTACCACCCCCAGTTCAATTGAGAGAGGGAAGCTGTCTGCTTCAAATCTATACTTGATTCCCCTTCCACTCTTTCTCT
CGCCTAGTTCTGATGATATGCAGGTTTTGGTGGAAAGAATCTGTAGCTTTTCAGTTTGAATCAGCACTCTTCTTGAGGACCTCCTA
TGTGTGCAGGACCATGGGATGTTGGAGAGAAGGGTATGACATGCATTCTTTTCTGGTAATTTCTTTTGGGCAAATCAGATACAGAC
TCTGTGTCTGTATCTTCAAGAAATGTAAGTCTAATGGAAAACAGACATAAGCTTGGCAAGCTT

10. GROW1_R

GGTACCTAAGGTACCTAAGCAGTAGTTTAGTAAGTGCACACCATATCCTGGGCACTTTGTGGGAACTAGGGAATAGTTCTCAAGAA
GCTCACAGTCTAGTGAGGGAGATAAGATGTGTAAAATGGTCAATTATAAATCAGAATGCCTAAGTTATATAATAAATATGAGTAAG
AAGTACTTATAGGATCACAAAGGAAGCTGCAACTATCAAAACCCTTATACTTCTACCTGTTAGTTCTTTGAAATGTGCCACGCATC
AGGTAGCTTAAAAGGCACACTGCCTGCCATCTTGTTGTCTGAATGGTGAAGGGTAAGGCATAGCGTTCAGGGAAGGAAGCTGCATA
TATTGAAGGGAAAAAGTGCTTTTTTAGGAGAGGAGGACTGTAGAGGTGGCACTGTAAAATTACTCACTGGAAAGGTGGGAGGGTTG
CAAGAAGGCTAGGGGAACTCAAAGATAATATTCAGAACCTGTATGAAAGACTCTTTATTGCCCATATAGCTAGAGGTATCAGTGTT
AGAGGGAGTGGATGGCAAAATAAAATAAAATATATAATCTCTGTGACCACATCCCAAGGTCAATACCTGACTAACAGCAGATGGGA
GAACCTTGAAACTACTTATAGTTCTTTAAAGTTTCCAAACTGTGCCTGCTTAATCTTCCTTGCTGAGGTTGGTTTAATTTCTGAGG
AGTAGGGTGTACCACCCCCAGTTCAATTGAGAGAGGGAAGCTGTCTGCTTCAAATCTATACTTGATTCCCCTTCCACTCTTTCTCT
CGCCTAGTTCTGATGATATGCAGGTTTTGGTGGAAAGAATCTGTAGCTTTTCAGTTTGAATCAGCACTCTTCTTGAGGACCTCCTA
TGTGTGCAGGACCATGGGATGTTGGAGAGAAGGGTATGACATGCATTCTTTTCTGGTAATTTCTTTTGGGCAAATCAGATACAGAC
TCTGTGTCTGTATCTTCAAGAAATGTAAGTCTAATGGAAAACAGACATAAGCTTGGCAAGCTT

11. 5UTR NR-NR

GGTACCGGATTCAAAACTAGGGGGAAAAAAAAACTGGAGCACACAGGCAGCATTACGCCATTCTTCCTTCTTGGAAAAATCCCTCA
GCCTTATACAAGCCTCCTTCAAGCCCTCAGTCAGTTGTGCAGGAGAAAGGGGGCGGTCGGCTTTCTCCTTTCAAGAACGAGTTATT
TTCAGCTGCTGACTGGAGACGGTGCACGTCTGGATACGAGAGCATTTCCACTATGGGACTGGATACAAACACACACCCGGCAGACT
TCAAGAGTCTCAGACTGAGGAGAAAGCCTTTCCTTCTGCTGCTACTGCTGCTGCCGCTGCTTTTGAAAGTCCACTCCTTTCATGGT
TTTTCCTGCCAAACCAGAGGCACCTTCGCTGCTGCCGCTGTTCTCTTTGGTGTCATTCAGCGGAAGCTT

12. 5UTR R-R

GGTACCGGATTCAAAACTAGGGGGAAAAAAAAACTGGAGCACACAGGCAGCATTACGCCATTCTTCCTTCTTGGAAAAATCCCTCA
GCCTTATACAAGCCTCCTTCAAGCCCTCAGTCAGTTGTGCAGGAGAAAGGGGGCGGTTGGCTTTCTCCTTTCAAGAACGAGTTATT
TTCAGCTGCTGACTGGAGACGGTGCACGTCTGGATACGAGAGCATTTCCACTATGGGACTGGATACAAACACACACCCGGCAGACT
TCAAGAGTCTCAGACTGAGGAGAAAGCCTTTCCTTCTGCTGCTACTGCTGCTGCCGCTGCTTTTGAAAGTCCACTCCTTTCATGGT
TTTTCCTGCCAAACCAGAGGCACCTTTGCTGCTGCCGCTGTTCTCTTTGGTGTCATTCAGCGGAAGCTT

13. 5UTR R-NR

GGTACCGGATTCAAAACTAGGGGGAAAAAAAAACTGGAGCACACAGGCAGCATTACGCCATTCTTCCTTCTTGGAAAAATCCCTCA
GCCTTATACAAGCCTCCTTCAAGCCCTCAGTCAGTTGTGCAGGAGAAAGGGGGCGGTTGGCTTTCTCCTTTCAAGAACGAGTTATT
TTCAGCTGCTGACTGGAGACGGTGCACGTCTGGATACGAGAGCATTTCCACTATGGGACTGGATACAAACACACACCCGGCAGACT
TCAAGAGTCTCAGACTGAGGAGAAAGCCTTTCCTTCTGCTGCTACTGCTGCTGCCGCTGCTTTTGAAAGTCCACTCCTTTCATGGT
TTTTCCTGCCAAACCAGAGGCACCTTCGCTGCTGCCGCTGTTCTCTTTGGTGTCATTCAGCGGAAGCTT

14. 5UTR NR-R

GGTACCGGATTCAAAACTAGGGGGAAAAAAAAACTGGAGCACACAGGCAGCATTACGCCATTCTTCCTTCTTGGAAAAATCCCTCA
GCCTTATACAAGCCTCCTTCAAGCCCTCAGTCAGTTGTGCAGGAGAAAGGGGGCGGTCGGCTTTCTCCTTTCAAGAACGAGTTATT
TTCAGCTGCTGACTGGAGACGGTGCACGTCTGGATACGAGAGCATTTCCACTATGGGACTGGATACAAACACACACCCGGCAGACT
TCAAGAGTCTCAGACTGAGGAGAAAGCCTTTCCTTCTGCTGCTACTGCTGCTGCCGCTGCTTTTGAAAGTCCACTCCTTTCATGGT
TTTTCCTGCCAAACCAGAGGCACCTTTGCTGCTGCCGCTGTTCTCTTTGGTGTCATTCAGCGGAAGCTT

15. R4 5UTR NR-NR-NR

GGTACCGATCTCCCATCAGTCCCTGTATCCACATAGTCTGGTTCCTCTGTTAAGCAGTCACATAAATATCTTTAAGGCAGCAACAG
TGAAAAGGGGAAAGAGAGCAGCGAATTGGCCTCAAATGAACTGGCATCTCTTTAAAGCTAATATTCCTCCTTGAGAATGCTACTTT
AATTAAACATGTGCCCTGCAAGATTCATTATTGCCTTACTTGTGTTCTCTGTTGGTTGTCTCATTCATTAATCTGCCGATTAGTAA

CCCTGAACTCTTGCATCCCTTCCTTTTTTTTGCATTCCCTTTTTAATATACTATGCTGTTTCCCCTGGGACATAAACTCCAGGACC
CGTCCCTGGCCTGCTCGTGTTTGTTTTCTTTGCCCCTTGTGCCTGCCTCTGCTATAATATTTGTAGTATGTGTTAAGTAATCCTTA
AGGTAACATTTTTATTATGTCATAAAAAGTGCATGGCAAATATATTATAACTCATGGACTTTGTTTAAAAATGAACGGATAATTTA
TTGAGTGTGTTACATGAGCCTCACTCCCCTCCCCATTTCAGCTGAGACCTGCTGCTCATTTAGCCGAGCCTCCTATTGAGTGCTCC
CTCCCTGTGACATGCTGGGAAAGCAGGCCTGTGGTTGGCTAGAGGTGACATCATAGTTTATGTACTCAATAAAAATTTAATTAAGT
CATTACAGGAATTGCAGTCATCATTTAACCCTCTGATTTCTTCTGTTTAGTTTATTATTAAACATTCAGTAATACCAGTCACTGAA
TCCTGGTGGCTTATAAAATATCTGCATGAATTTTAATGGCTTAAATAAATGGGAAAATATGAGTAACTGATATAAAAGGAACCTTT
TCCTTGTGGTGTTGTATGTATGGGTTTTTATGAAAAGCAAAGCATTATGGAAAGGATGAGTTTAATAAGAGAACGGCCCAACACTA
TCTGGGAGGAGTTTCTCTTCTGCTGGGTGCAGCTGTGTAGTCATGGGCAGAGTGCTCCATATAGAATGGTTTAAACGGATTCAAAA
CTAGGGGGAAAAAAAAACTGGAGCACACAGGCAGCATTACGCCATTCTTCCTTCTTGGAAAAATCCCTCAGCCTTATACAAGCCTC
CTTCAAGCCCTCAGTCAGTTGTGCAGGAGAAAGGGGGCGGTCGGCTTTCTCCTTTCAAGAACGAGTTATTTTCAGCTGCTGACTGG
AGACGGTGCACGTCTGGATACGAGAGCATTTCCACTATGGGACTGGATACAAACACACACCCGGCAGACTTCAAGAGTCTCAGACT
GAGGAGAAAGCCTTTCCTTCTGCTGCTACTGCTGCTGCCGCTGCTTTTGAAAGTCCACTCCTTTCATGGTTTTTCCTGCCAAACCA
GAGGCACCTTCGCTGCTGCCGCTGTTCTCTTTGGTGTCATTCAGCGGAAGCTT

16. R4 5UTR NR-NR-R:
GGTACCGATCTCCCATCAGTCCCTGTATCCACATAGTCTGGTTCCTCTGTTAAGCAGTCACATAAATATCTTTAAGGCAGCAACAG
TGAAAAGGGGAAAGAGAGCAGCGAATTGGCCTCAAATGAACTGGCATCTCTTTAAAGCTAATATTCCTCCTTGAGAATGCTACTTT
AATTAAACATGTGCCCTGCAAGATTCATTATTGCCTTACTTGTGTTCTCTGTTGGTTGTCTCATTCATTAATCTGCCGATTAGTAA
CCCTGAACTCTTGCATCCCTTCCTTTTTTTTGCATTCCCTTTTTAATATACTATGCTGTTTCCCCTGGGACATAAACTCCAGGACC
CGTCCCTGGCCTGCTCGTGTTTGTTTTCTTTGCCCCTTGTGCCTGCCTCTGCTATAATATTTGTAGTATGTGTTAAGTAATCCTTA
AGGTAACATTTTTATTATGTCATAAAAAGTGCATGGCAAATATATTATAACTCATGGACTTTGTTTAAAAATGAACGGATAATTTA
TTGAGTGTGTTACATGAGCCTCACTCCCCTCCCCATTTCAGCTGAGACCTGCTGCTCATTTAGCCG
AGCCTCCTATTGAGTGCTCCCTCCCTGTGACATGCTGGGAAAGCAGGCCTGTGGTTGGCTAGAGGTGACATCATAGTTTATGTACT
CAATAAAAATTTAATTAAGTCATTACAGGAATTGCAGTCATCATTTAACCCTCTGATTTCTTCTGTTTAGTTTATTATTAAACATT
CAGTAATACCAGTCACTGAATCCTGGTGGCTTATAAAATATCTGCATGAATTTTAATGGCTTAAATAAATGGGAAAATATGAGTAA
CTGATATAAAAGGAACCTTTTCCTTGTGGTGTTGTATGTATGGGTTTTTATGAAAAGCAAAGCATTATGGAAAGGATGAGTTTAAT
AAGAGAACGGCCCAACACTATCTGGGAGGAGTTTCTCTTCTGCTGGGTGCAGCTGTGTAGTCATGGGCAGAGTGCTCCATATAGAA
TGGTTTAAACGGATTCAAAACTAGGGGGAAAAAAAAACTGGAGCACACAGGCAGCATTACGCCATTCTTCCTTCTTGGAAAAATCC
CTCAGCCTTATACAAGCCTCCTTCAAGCCCTCAGTCAGTTGTGCAGGAGAAAGGGGGCGGTCGGCTTTCTCCTTTCAAGAACGAGT
TATTTTCAGCTGCTGACTGGAGACGGTGCACGTCTGGATACGAGAGCATTTCCACTATGGGACTGGATACAAACACACACCCGGCA
GACTTCAAGAGTCTCAGACTGAGGAGAAAGCCTTTCCTTCTGCTGCTACTGCTGCTGCCGCTGCTTTTGAAAGTCCACTCCTTTCA
TGGTTTTTCCTGCCAAACCAGAGGCACCTTTGCTGCTGCCGCTGTTCTCTTTGGTGTCATTCAGCGGAAGCTT

17. R4 5UTR NR-R-NR
GGTACCGATCTCCCATCAGTCCCTGTATCCACATAGTCTGGTTCCTCTGTTAAGCAGTCACATAAATATCTTTAAGGCAGCAACAG
TGAAAAGGGGAAAGAGAGCAGCGAATTGGCCTCAAATGAACTGGCATCTCTTTAAAGCTAATATTCCTCCTTGAGAATGCTACTTT
AATTAAACATGTGCCCTGCAAGATTCATTATTGCCTTACTTGTGTTCTCTGTTGGTTGTCTCATTCATTAATCTGCCGATTAGTAA
CCCTGAACTCTTGCATCCCTTCCTTTTTTTTGCATTCCCTTTTTAATATACTATGCTGTTTCCCCTGGGACATAAACTCCAGGACC
CGTCCCTGGCCTGCTCGTGTTTGTTTTCTTTGCCCCTTGTGCCTGCCTCTGCTATAATATTTGTAGTATGTGTTAAGTAATCCTTA
AGGTAACATTTTTATTATGTCATAAAAAGTGCATGGCAAATATATTATAACTCATGGACTTTGTTTAAAAATGAACGGATAATTTA
TTGAGTGTGTTACATGAGCCTCACTCCCCTCCCCATTTCAGCTGAGACCTGCTGCTCATTTAGCCGAGCCTCCTATTGAGTGCTCC
CTCCCTGTGACATGCTGGGAAAGCAGGCCTGTGGTTGGCTAGAGGTGACATCATAGTTTATGTACTCAATAAAAATTTAATTAAGT
CATTACAGGAATTGCAGTCATCATTTAACCCTCTGATTTCTTCTGTTTAGTTTATTATTAAACATTCAGTAATACCAGTCACTGAA
TCCTGGTGGCTTATAAAATATCTGCATGAATTTTAATGGCTTAAATAAATGGGAAAATATGAGTAACTGATATAAAAGGAACCTTT
TCCTTGTGGTGTTGTATGTATGGGTTTTTATGAAAAGCAAAGCATTATGGAAAGGATGAGTTTAATAAGAGAACGGCCCAACACTA
TCTGGGAGGAGTTTCTCTTCTGCTGGGTGCAGCTGTGTAGTCATGGGCAGAGTGCTCCATATAGAATGGTTTAAACGGATTCAAAA
CTAGGGGGAAAAAAAAACTGGAGCACACAGGCAGCATTACGCCATTCTTCCTTCTTGGAAAAATCCCTCAGCCTTATACAAGCCTC
CTTCAAGCCCTCAGTCAGTTGTGCAGGAGAAAGGGGGCGGTTGGCTTTCTCCTTTCAAGAACGAGTTATTTTCAGCTGCTGACTGG
AGACGGTGCACGTCTGGATACGAGAGCATTTCCACTATGGGACTGGATACAAACACACACCCGGCAGACTTCAAGAGTCTCAGACT
GAGGAGAAAGCCTTTCCTTCTGCTGCTACTGCTGCTGCCGCTGCTTTTGAAAGTCCACTCCTTTCATGGTTTTTCCTGCCAAACCA
GAGGCACCTTCGCTGCTGCCGCTGTTCTCTTTGGTGTCATTCAGCGGAAGCTT

18. R4 5UTR NR-R-R
GGTACCGATCTCCCATCAGTCCCTGTATCCACATAGTCTGGTTCCTCTGTTAAGCAGTCACATAAATATCTTTAAGGCAGCAACAG
TGAAAAGGGGAAAGAGAGCAGCGAATTGGCCTCAAATGAACTGGCATCTCTTTAAAGCTAATATTCCTCCTTGAGAATGCTACTTT
AATTAAACATGTGCCCTGCAAGATTCATTATTGCCTTACTTGTGTTCTCTGTTGGTTGTCTCATTCATTAATCTGCCGATTAGTAA
CCCTGAACTCTTGCATCCCTTCCTTTTTTTTGCATTCCCTTTTTAATATACTATGCTGTTTCCCCTGGGACATAAACTCCAGGACC
CGTCCCTGGCCTGCTCGTGTTTGTTTTCTTTGCCCCTTGTGCCTGCCTCTGCTATAATATTTGTAGTATGTGTTAAGTAATCCTTA
AGGTAACATTTTTATTATGTCATAAAAAGTGCATGGCAAATATATTATAACTCATGGACTTTGTTTAAAAATGAACGGATAATTTA
TTGAGTGTGTTACATGAGCCTCACTCCCCTCCCCATTTCAGCTGAGACCTGCTGCTCATTTAGCCGAGCCTCCTATTGAGTGCTCC
CTCCCTGTGACATGCTGGGAAAGCAGGCCTGTGGTTGGCTAGAGGTGACATCATAGTTTATGTACTCAATAAAAATTTAATTAAGT

CATTACAGGAATTGCAGTCATCATTTAACCCTCTGATTTCTTCTGTTTAGTTTATTATTAAACATTCAGTAATACCAGTCACTGAA
TCCTGGTGGCTTATAAAATATCTGCATGAATTTTAATGGCTTAAATAAATGGGAAAATATGAGTAACTGATATAAAAGGAACCTTT
TCCTTGTGGTGTTGTATGTATGGGTTTTTATGAAAAGCAAAGCATTATGGAAAGGATGAGTTTAATAAGAGAACGGCCCAACACTA
TCTGGGAGGAGTTTCTCTTCTGCTGGGTGCAGCTGTGTAGTCATGGGCAGAGTGCTCCATATAGAATGGTTTAAACGGATTCAAAA
CTAGGGGGAAAAAAAAACTGGAGCACACAGGCAGCATTACGCCATTCTTCCTTCTTGGAAAAATCCCTCAGCCTTATACAAGCCTC
CTTCAAGCCCTCAGTCAGTTGTGCAGGAGAAAGGGGGCGGTTGGCTTTCTCCTTTCAAGAACGAGTTATTTTCAGCTGCTGACTGG
AGACGGTGCACGTCTGGATACGAGAGCATTTCCACTATGGGACTGGATACAAACACACACCCGGCAGACTTCAAGAGTCTCAGACT
GAGGAGAAAGCCTTTCCTTCTGCTGCTACTGCTGCTGCCGCTGCTTTTGAAAGTCCACTCCTTTCATGGTTTTTCCTGCCAAACCA
GAGGCACCTTTGCTGCTGCCGCTGTTCTCTTTGGTGTCATTCAGCGGAAGCTT

19. R4 5UTR R-NR-NR
GGTACCGATCTCCCATCAGTCCCTGTATCCACATAGTCTGGTTCCTCTGTTAAGCAGTCACATAAATATCTTTAAGGCAGCAACAG
TGAAAAGGGGAAAGAGAGCAGCGAATTGGCCTCAAATGAACTGGCATCTCTTTAAAGCTAATATTCCTCCTTGAGAATGCTACTTT
AATTAAACATGTGCCCTGCAAATTCATTATTGCCTTACTTGTGTTCTCTGTTGGTTGTCTCATTCATTAATCTGCCGATTAGTAAC
CCTGAACTCTTGCATCCCTTCCTTTTTTTTGCATTCCCTTTTTAATATACTATGCTGTTTCCCCTGGGACATAAACTCCAGGACCC
GTCCCTGGCCTGCTCGTGTTTGTTTTCTTTGCCCCTTGTGCCTGCCTCTGCTATAATATTTGTAGTATGTGTTAAGTAATCCTTAA
GGTAACATTTTTATTATGTCATAAAAAGTGCATGGCAAATATATTATAACTCATGGACTTTGTTTAAAAATGAACGGATAATTTAT
TGAGTGTGTTACATGAGCCTCACTCCCCTCCCCATTTCAGCTGAGACCTGCTGCTCATTTAGCCAAGCCTCCTATTGAGTGCTCCC
TCCCTGTGACATGCTGGGAAAGCAGGCCTGTGGTTGGCTAGAGGTGACATCATAGTTTATGTACTCAATAAAAATTTAATTAAGTC
ATTACAGGAATTGCAGTCATCATTTAACCCTCTGATTTCTTCTGTTTAGTTTATTATTAAACATTCAGTAATACCAGTCACTGAAT
CCTGGTGGCTTATAAAATATCTGCATGAATTTTAATGGCTTAAATAAATGGGAAAATATGAGTAACTGATATAAAAGGAACCTTTT
CCTTGTGGTGTTGTATGTATGGGTTTTTATGAAAAGCAAAGCATTATGGAAAGGATGAGTTTAATAAGAGAACGGCCCAACACTAT
CTGGGAGGAGTTTCTCTTCTGCTGGGTGCAGCTGTGTAGTCATGGGCAGAGTGCTCCATATAGAATGGTTTAAACGGATTCAAAAC
TAGGGGGAAAAAAAAACTGGAGCACACAGGCAGCATTACGCCATTCTTCCTTCTTGGAAAAATCCCTCAGCCTTATACAAGCCTCC
TTCAAGCCCTCAGTCAGTTGTGCAGGAGAAAGGGGGCGGTCGGCTTTCTCCTTTCAAGAACGAGTTATTTTCAGCTGCTGACTGGA
GACGGTGCACGTCTGGATACGAGAGCATTTCCACTATGGGACTGGATACAAACACACACCCGGCAGACTTCAAGAGTCTCAGACTG
AGGAGAAAGCCTTTCCTTCTGCTGCTACTGCTGCTGCCGCTGCTTTTGAAAGTCCACTCCTTTCATGGTTTTTCCTGCCAAACCAG
AGGCACCTTCGCTGCTGCCGCTGTTCTCTTTGGTGTCATTCAGCGGAAGCTT

20. R4 5UTR R-NR-R
GGTACCGATCTCCCATCAGTCCCTGTATCCACATAGTCTGGTTCCTCTGTTAAGCAGTCACATAAATATCTTTAAGGCAGCAACAG
TGAAAAGGGGAAAGAGAGCAGCGAATTGGCCTCAAATGAACTGGCATCTCTTTAAAGCTAATATTCCTCCTTGAGAATGCTACTTT
AATTAAACATGTGCCCTGCAAGATTCATTATTGCCTTACTTGTGTTCTCTGTTGGTTGTCTCATTCATTAATCTGCCGATTAGTAA
CCCTGAACTCTTGCATCCCTTCCTTTTTTTTGCATTCCCTTTTTAATATACTATGCTGTTTCCCCTGGGACATAAACTCCAGGACC
CGTCCCTGGCCTGCTCGTGTTTGTTTTCTTTGCCCCTTGTGCCTGCCTCTGCTATAATATTTGTAGTATGTGTTAAGTAATCCTTA
AGGTAACATTTTTATTATGTCATAAAAAGTGCATGGCAAATATATTATAACTCATGGACTTTGTTTAAAAATGAACGGATAATTTA
TTGAGTGTGTTACATGAGCCTCACTCCCCTCCCCATTTCAGCTGAGACCTGCTGCTCATTTAGCCAAGCCTCCTATTGAGTGCTCC
CTCCCTGTGACATGCTGGGAAAGCAGGCCTGTGGTTGGCTAGAGGTGACATCATAGTTTATGTACTCAATAAAAATTTAATTAAGT
CATTACAGGAATTGCAGTCATCATTTAACCCTCTGATTTCTTCTGTTTAGTTTATTATTAAACATTCAGTAATACCAGTCACTGAA
TCCTGGTGGCTTATAAAATATCTGCATGAATTTTAATGGCTTAAATAAATGGGAAAATATGAGTAACTGATATAAAAGGAACCTTT
TCCTTGTGGTGTTGTATGTATGGGTTTTTATGAAAAGCAAAGCATTATGGAAAGGATGAGTTTAATAAGAGAACGGCCCAACACTA
TCTGGGAGGAGTTTCTCTTCTGCTGGGTGCAGCTGTGTAGTCATGGGCAGAGTGCTCCATATAGAATGGTTTAAACGGATTCAAAA
CTAGGGGGAAAAAAAAACTGGAGCACACAGGCAGCATTACGCCATTCTTCCTTCTTGGAAAAATCCCTCAGCCTTATACAAGCCTC
CTTCAAGCCCTCAGTCAGTTGTGCAGGAGAAAGGGGGCGGTCGGCTTTCTCCTTTCAAGAACGAGTTATTTTCAGCTGCTGACTGG
AGACGGTGCACGTCTGGATACGAGAGCATTTCCACTATGGGACTGGATACAAACACACACCCGGCAGACTTCAAGAGTCTCAGACT
GAGGAGAAAGCCTTTCCTTCTGCTGCTACTGCTGCTGCCGCTGCTTTTGAAAGTCCACTCCTTTCATGGTTTTTCCTGCCAAACCA
GAGGCACCTTTGCTGCTGCCGCTGTTCTCTTTGGTGTCATTCAGCGGAAGCTT

21. R4 5UTR R-R-NR
GGTACCGATCTCCCATCAGTCCCTGTATCCACATAGTCTGGTTCCTCTGTTAAGCAGTCACATAAATATCTTTAAGGCAGCAACAG
TGAAAAGGGGAAAGAGAGCAGCGAATTGGCCTCAAATGAACTGGCATCTCTTTAAAGCTAATATTCCTCCTTGAGAATGCTACTTT
AATTAAACATGTGCCCTGCAAGATTCATTATTGCCTTACTTGTGTTCTCTGTTGGTTGTCTCATTCATTAATCTGCCGATTAGTAA
CCCTGAACTCTTGCATCCCTTCCTTTTTTTTGCATTCCCTTTTTAATATACTATGCTGTTTCCCCTGGGACATAAACTCCAGGACC
CGTCCCTGGCCTGCTCGTGTTTGTTTTCTTTGCCCCTTGTGCCTGCCTCTGCTATAATATTTGTAGTATGTGTTAAGTAATCCTTA
AGGTAACATTTTTATTATGTCATAAAAAGTGCATGGCAAATATATTATAACTCATGGACTTTGTTTAAAAATGAACGGATAATTTA
TTGAGTGTGTTACATGAGCCTCACTCCCCTCCCCATTTCAGCTGAGACCTGCTGCTCATTTAGCCAAGCCTCCTATTGAGTGCTCC
CTCCCTGTGACATGCTGGGAAAGCAGGCCTGTGGTTGGCTAGAGGTGACATCATAGTTTATGTACTCAATAAAAATTTAATTAAGT
CATTACAGGAATTGCAGTCATCATTTAACCCTCTGATTTCTTCTGTTTAGTTTATTATTAAACATTCAGTAATACCAGTCACTGAA
TCCTGGTGGCTTATAAAATATCTGCATGAATTTTAATGGCTTAAATAAATGGGAAAATATGAGTAACTGATATAAAAGGAACCTTT
TCCTTGTGGTGTTGTATGTATGGGTTTTTATGAAAAGCAAAGCATTATGGAAAGGATGAGTTTAATAAGAGAACGGCCCAACACTA
TCTGGGAGGAGTTTCTCTTCTGCTGGGTGCAGCTGTGTAGTCATGGGCAGAGTGCTCCATATAGAATGGTTTAAACGGATTCAAAA
CTAGGGGGAAAAAAAAACTGGAGCACACAGGCAGCATTACGCCATTCTTCCTTCTTGGAAAAATCCCTCAGCCTTATACAAGCCTC

CTTCAAGCCCTCAGTCAGTTGTGCAGGAGAAAGGGGGCGGTTGGCTTTCTCCTTTCAAGAACGAGTTATTTTCAGCTGCTGACTGG
AGACGGTGCACGTCTGGATACGAGAGCATTTCCACTATGGGACTGGATACAAACACACACCCGGCAGACTTCAAGAGTCTCAGACT
GAGGAGAAAGCCTTTCCTTCTGCTGCTACTGCTGCTGCCGCTGCTTTTGAAAGTCCACTCCTTTCATGGTTTTTCCTGCCAAACCA
GAGGCACCTTCGCTGCTGCCGCTGTTCTCTTTGGTGTCATTCAGCGGAAGCTT

22. R4 5UTR R-R-R
GGTACCGATCTCCCATCAGTCCCTGTATCCACATAGTCTGGTTCCTCTGTTAAGCAGTCACATAAATATCTTTAAGGCAGCAACAG
TGAAAAGGGGAAAGAGAGCAGCGAATTGGCCTCAAATGAACTGGCATCTCTTTAAAGCTAATATTCCTCCTTGAGAATGCTACTTT
AATTAAACATGTGCCCTGCAAGATTCATTATTGCCTTACTTGTGTTCTCTGTTGGTTGTCTCATTCATTAATCTGCCGATTAGTAA
CCCTGAACTCTTGCATCCCTTCCTTTTTTTTGCATTCCCTTTTTAATATACTATGCTGTTTCCCCTGGGACATAAACTCCAGGACC
CGTCCCTGGCCTGCTCGTGTTTGTTTTCTTTGCCCCTTGTGCCTGCCTCTGCTATAATATTTGTAGTATGTGTTAAGTAATCCTTA
AGGTAACATTTTTATTATGTCATAAAAAGTGCATGGCAAATATATTATAACTCATGGACTTTGTTTAAAAATGAACGGATAATTTA
TTGAGTGTGTTACATGAGCCTCACTCCCCTCCCCATTTCAGCTGAGACCTGCTGCTCATTTAGCCAAGCCTCCTATTGAGTGCTCC
CTCCCTGTGACATGCTGGGAAAGCAGGCCTGTGGTTGGCTAGAGGTGACATCATAGTTTATGTACTCAATAAAAATTTAATTAAGT
CATTACAGGAATTGCAGTCATCATTTAACCCTCTGATTTCTTCTGTTTAGTTTATTATTAAACATTCAGTAATACCAGTCACTGAA
TCCTGGTGGCTTATAAAATATCTGCATGAATTTTAATGGCTTAAATAAATGGGAAAATATGAGTAACTGATATAAAAGGAACCTTT
TCCTTGTGGTGTTGTATGTATGGGTTTTTATGAAAAGCAAAGCATTATGGAAAGGATGAGTTTAATAAGAGAACGGCCCAACACTA
TCTGGGAGGAGTTTCTCTTCTGCTGGGTGCAGCTGTGTAGTCATGGGCAGAGTGCTCCATATAGAATGGTTTAAACGGATTCAAAA
CTAGGGGGAAAAAAAAACTGGAGCACACAGGCAGCATTACGCCATTCTTCCTTCTTGGAAAAATCCCTCAGCCTTATACAAGCCTC
CTTCAAGCCCTCAGTCAGTTGTGCAGGAGAAAGGGGGCGGTTGGCTTTCTCCTTTCAAGAACGAGTTATTTTCAGCTGCTGACTGG
AGACGGTGCACGTCTGGATACGAGAGCATTTCCACTATGGGACTGGATACAAACACACACCCGGCAGACTTCAAGAGTCTCAGACT
GAGGAGAAAGCCTTTCCTTCTGCTGCTACTGCTGCTGCCGCTGCTTTTGAAAGTCCACTCCTTTCATGGTTTTTCCTGCCAAACCA
GAGGCACCTTTGCTGCTGCCGCTGTTCTCTTTGGTGTCATTCAGCGGAAGCTT

**CRISPR-Cas9 single guide RNA sequences and locations**

Deletion of *R9* pertaining to Figure 2d:
Single guide L4: CACCGCGAGTTCCCAAGCGATG|CTG cut at hg19 chr20:33,925,323. Single guide R5: CACCGTCTTGGTCTTTGGCAGT|CCT cut at hg19 chr20:33,926,134. Relative luciferase expression of plasmid constructs targeting different combinations or risk and non-risk variants at rs2378349 (non-risk "A", risk "C") and rs2248393 (risk "C", non-risk "G") within regulatory region *R9*; chr20:33925385-33925404 (hg19) 3'(Hind III) GCG AAG CTT CTT TCC AGT TTA AAC AGT CT and; chr20:33926089-33926109 (hg19) 5'(Kpn1) CGC GGT ACC TTT CTT GAA TCT AAA GGC ATT.

Testing effects of *R2de* deletion (single guides below) in human T/C28a2 cells on *UQCC*, *GDF5* and *CEP250* expression:
Single guide L1: CACCGCTTGAAGGAGGCTTGT|ATA (cut at hg19 chr20:34,026,033)
Single guide 2: CACCGGAGGCTTGTATAAGGC|TGA (cut at hg19 chr20:34,026,040)
Single guide R6: CAGCGGGACTCTAGGCT|TAG (cut at hg19 chr20:34,026,401)
Single guide R9: CACCGAGCGGGACTCTAGGCTT|AGG (cut at hg19 chr20:34,026,402)

| Construct | Mm9/Hg19 coordinates or construct number | Region size | Tested (E) | # LacZ embryos | # reproducible (pattern) | Database or reference |
|---|---|---|---|---|---|---|
| 37 Kb region | hg19 ABC82134240H3 | 37 kb | E14.5 | 4 | 3 (sub-perichondral) | Human Structural Variation Project; Capellini et al., 2017; Nature Genetics |
| PHC17 | mm9 chr2:155,717,545-155,729,712 | 12.17 kb | E14.5 | 19 | 12 (entire growth plate) | Capellini et al., 2017; Nature Genetics |
| GROW1 | hg19 chr20:33,952,180-33,954,726 | 2.54 kb | E14.5 | 8 | 7 (sub-perichondral) | Capellini et al., 2017; Nature Genetics |
| R18-20 (PHC26) | mm9 chr2:155,720,654-155,722,130 | 1477 bp | E14.5 | 12 | 7 (humerus/shoulder) | New |
| R7 | mm9 chr2:155,717,572-155,717,882 | 311 bp | E14.5 | 11 | 7 (digits) | New |
| PHC18 | mm9 chr2:155,700,028-155,710,095 | ~10 kb | E14.5 | 18 | 11 (shoulder) | New |
| R8 | mm9 chr2:155,708,447-155,709,489 | 1043 bp | E14.5 | 6 | 4 (elbow,shoulder,knee, digits) | New |
| R9 | mm9 chr2:155,702,138-155,702,803 | 666 bp | E14.5 | 10 | 1 (wrist) | New |
| 41 kb region | hg19 ABC1246947700I24 | 41 kb | E14.5 | 4 | 3 (joints) | Human Structural Variation Project; Capellini et al., 2017; Nature Genetics |
| PHC19 | various, concatonated | 1898 bp | E14.5 | 7 | 7 (joints) | Chen, Capellini et al., 2016; Plos Genetics |
| R3 | see reference | 586 bp | E14.5 | 4 | 4 (interdigital webbing, joints) | Chen, Capellini et al., 2016; Plos Genetics |
| R4 | see reference | 975 bp | E14.5 | 11 | 11 (joints) | Chen, Capellini et al., 2016; Plos Genetics |
| R5 | see reference | 337 bp | E14.5 | 13 | 8 (entire digit (prechondrogenic mesenchyme) | Chen, Capellini et al., 2016; Plos Genetics |
| R3+R4 | various, concatonated | 1561 bp | E14.5 | 14 | 9 (shoulder, elbow, knee, digits, webbing) | New |
| R3+R5 | various, concatonated | 923 bp | E14.5 | 2 | 2 (webbing) | New |
| R4+R5 | various, concatonated | 1312 bp | E14.5 | 12 | digits) | New |
| R4, All MUTB sites | various, concatonated | 1898 bp | E14.5 | 8 | 4 (digit stripes, lost knee and elbow in all) | New |
| R4, MUTB1 | various, concatonated | 1898 bp | E14.5 | 17 | elbow) | New |
| R4, MUTB2 | various, concatonated | 1898 bp | E14.5 | 8 | 8 (digits, webbing; slight reduction in expression in knee and elbow) | New |
| R4, MUTB3 | various, concatonated | 1898 bp | E14.5 | 9 | and elbow in 1) | New |
| R4, MUTB4 | various, concatonated | 1898 bp | E14.5 | 18 | 16 (digits, webbing, knee, elbow) | New |
| R4, MUTB5 | various, concatonated | 1898 bp | E14.5 | 11 | 9 (digits, webbing, knee, elbow) | New |
| R2a-e | see reference | 802 bp | E14.5 | 9 | 8 (joints) | Chen, Capellini et al., 2016; Plos Genetics |
| R2abc | see reference | 407 bp | E14.5 | 16 | 1 (no consisent expression) | Chen, Capellini et al., 2016; Plos Genetics |
| R2de | see reference | 395 bp | E14.5 | 5 | 4 (FL, HL joints) | Chen, Capellini et al., 2016; Plos Genetics |
| R2d | see reference | 215 bp | E14.5 | 9 | 6 (HL joints) | Chen, Capellini et al., 2016; Plos Genetics |
| R2e | see reference | 206 bp | E14.5 | 11 | 7 (FL joints) | Chen, Capellini et al., 2016; Plos Genetics |
| R2abc+d | various, concatonated | 622 bp | E14.5 | 16 | 5 (no consistent expression) | New |
| R2abc+e | various, concatonated | 613 bp | E14.5 | 4 | 1 (no consisent expression) | New |
| R2d103bp | within subregion d | 103 bp | E14.5 | 9 | 5 (HL joints) | New |

**Supplementary Table 1.** Summary of expression patterns generated by BACs or smaller regulatory or partial regulatory region constructs in independent E14.5 transgenic embryos.

| Construct | Purpose | Source DNA | Coordinates (hg19)/Cat.# | Forward (5' to 3') or sgRNA seq |
|---|---|---|---|---|
| pGL 4.23 Forward | Sequencing primer for insert region in pGL4.23 | pGL4.23 vector | pGL4.23 vector: 5146-5165 | GGCTGTCCCCAGTGCAAGTG |
| PGL 4.23 Reverse | Sequencing primer for insert region in pGL4.23 | pGL4.23 vector | pGL4.23 vector: 1129-1148 | CAGGGCGTAGCGCTTCATGG |
| R9_HindIII_For | Cloning of R9 in luciferase vector (HindIII linker) | human samples | chr20:33925385-33925404 | GCG AAG CTT CTT TCC AGT TTA AAC AGT CT |
| R9_Kpn1_Rev | Cloning of R9 in luciferase vector (KpnI linker) | human samples | chr20:33926089-33926109 | CGC GGT ACC TTT CTT GAA TCT AAA GGC ATT |
| R9_HindIII_For_2 | Cloning of R9 in luciferase vector (HindIII linker) | human samples | chr20:33925366-33925385 | GCC AAG CTT TCA GAT TAT TTC AGG GTT CC |
| R9_HindIII_For_2' | Cloning of R9 in luciferase vector (HindIII linker) | human samples | chr20:33925365-33925384 | GCC AAG CTT ATC AGA TTA TTT CAG GGT TC |
| *R2de* CRISPR L1 | Testing effects of *R2de* deletion | human samples | chr20:34,026,033 | CACCGCTTGAAGGAGGCTTGT\|ATA |
| R2de CRISPR L2 | Testing effects of *R2de* deletion | human samples | chr20:34,026,040 | CACCGGAGGCTTGTATAAGGC\|TGA |
| R2de CRISPR R6 | Testing effects of *R2de* deletion | human samples | chr20:34,026,401 | CAGCGGGACTCTAGGCT\|TAG |
| R2de CRISPR R9 | Testing effects of *R2de* deletion | human samples | chr20:34,026,402 | CACCGAGCGGGACTCTAGGCTT\|AG G |
| R4_HindIII_For | Cloning of R4 in luciferase vector (HindIII linker) | human samples | chr20:33906729-33906746 | ACC AAG CTT CAT TCT ATA TGG AGC ACT |
| R*4*_Kpn1_Rev | Cloning of R4 in luciferase vector (KpnII linker) | human samples | chr20:33907709-33907726 | TAA CCA TGG TCA GGG ACA TAG GTG TA |
| Rs6060369 CRISPR L1 | sgRNA targeting | Integrated DNA technologies | chr20:33907091-33907110 | CACCGTGTCACCTCTAGCCAACCAC |
| Rs6060369 CRISPR R1 | sgRNA targeting | Integrated DNA technologies | chr20:33907091-33907110 | AAACGTGGTTGGCTAGAGGTGACA C |
| Rs6060369 CRISPR L2 | sgRNA targeting | Integrated DNA technologies | chr20:33907136-33907155 | CACCGGGAGGGAGCACTCAATAGG |
| Rs6060369 CRISPR R2 | sgRNA targeting | Integrated DNA technologies | chr20:33907136-33907155 | AAACCCTATTGAGTGCTCCCTCCC |
| Rs6060369 CRISPR L3 | sgRNA targeting | Integrated DNA technologies | chr20:33907141-33907160 | CACCGGAGCACTCAATAGGAGGCT |
| Rs6060369 CRISPR R3 | sgRNA targeting | Integrated DNA technologies | chr20:33907141-33907160 | AAACAGCCTCCTATTGAGTGCTCC |
| Rs6060369 CRISPR L4 | sgRNA targeting | Integrated DNA technologies | chr20:33907157-33907166 | CACCGGCTTGGCTAAATGAGCAGC |
| Rs6060369 CRISPR R4 | sgRNA targeting | Integrated DNA technologies | chr20:33907157-33907166 | AAACGCTGCTCATTTAGCCAAGCC |
| Rs6060369 CRISPR L5 | sgRNA targeting | Integrated DNA technologies | chr20:33907115-33907134 | CACCGCTGCTTTCCCAGCATGTCAC |
| Rs6060369 CRISPR R5 | sgRNA targeting | Integrated DNA technologies | chr20:33907115-33907134 | AAACGTGACATGCTGGGAAAGCAGC |
| Rs6060369 CRISPR L7 | sgRNA targeting | Integrated DNA technologies | chr20:33907177-33907196 | CACCGAGGTCTCAGCTGAAATGGG G |
| Rs6060369 CRISPR R7 | sgRNA targeting | Integrated DNA technologies | chr20:33907177-33907196 | AAACCCCCATTTCAGCTGAGACCTC |
| human GADPH F | qPCR primer | Integrated DNA technologies | N/A | CACCGAAGGTGAAGGTCGGAGTC |
| human GADPH R | qPCR primer | Integrated DNA technologies | N/A | AAACGAAGATGGTGATGGGATTTC |

**Supplementary Table 2.** CRISPR guide sequences and locations and PCR sequences.
